# Biological Time Equivalence in Vertebrates: Thermodynamic Framework, Comparative Tests, and Clade-Specific Deviations

**DOI:** 10.64898/2026.03.27.714731

**Authors:** Mesfin Asfaw Taye

## Abstract

Across adult warm-blooded vertebrates, the product of resting heart rate *f*_H_ and maximum lifespan *L* is approximately constant: *N*_⋆_ = *f*_H_ *L* ≈ 10^9^ cardiac cycles. This empirical regularity, noted since Rubner (1908), has lacked a widely accepted thermodynamic interpretation. We derive *N*_⋆_ ≈ 10^9^ from the non-equilibrium second law by treating the adult organism as a metabolic non-equilibrium steady state (NESS) and introducing the empirical closure *ė*_p_ = *σ*_0_*f*, which links entropy production rate to heart rate via a mass-specific parameter *σ*_0_ ∝ *M*^0^. Under this closure, the lifetime entropy budget ∑ = *σ*_0_*N*_⋆_ is approximately species-independent when *σ*_0_ is approximately constant—a condition whose direct calorimetric verification remains the critical outstanding experimental test. We further show that *N*_⋆_ is the correct primitive invariant: lifetime energy per unit mass is a derived consequence, valid only when body temperature and the mass-specific entropy cost per cycle are both approximately constant. This framework, which we term the Principle of Biological Time Equivalence (PBTE), is placed on a fully falsifiable footing with explicit assumptions, a domain-of-validity table, and five numerical falsification criteria. We test the framework against a dataset of 230 adult vertebrate species spanning eight taxonomic groups. Ordinary least-squares regression on the *n* = 43 directly measured non-primate placentals yields slope 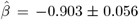 (*R*^2^ = 0.863; *F* -test *p* = 0.093 against *β* = −1). Phylogenetically independent contrasts on 112 endotherm species yield a log_10_ *f*_H_–log_10_ *L* slope of −0.99 ± 0.04 (*p* = 0.84 against slope −1), confirming the relation is not a phylogenetic artefact. The WBE kinematic null of zero inter-clade variation is rejected (*F* = 12.7, *p* < 0.001). Four warm-blooded clades depart systematically from the mammalian baseline; we derive their longevity deviations from a unified thermodynamic multiplier Φ_C_ = Φ_duty_ · Φ_thermal_ · Φ_mito+oxid_ · Φ_haz_, calibrated to independently measured physiology. For primates, the elevated count ⟨*N*_⋆_⟩ ≈ (2–3) × 10^9^ follows from a neuro-metabolic entropy model in which greater neural metabolic investment reduces entropy produced per cardiac cycle. For bats, the extreme longevity (Φ_bat_ ≈ 7.9) arises from the multiplicative synergy of cardiac suppression during torpor and an Arrhenius thermal factor during hibernation—two mechanisms acting simultaneously whose thermodynamic motivation has not previously been given. For birds, an adverse thermal penalty (Φ_thermal_ = 0.73) and adverse flight duty cycle (Φ_duty_ = 0.87) are overcome by mitochondrial coupling efficiency and antioxidant robustness. For cetaceans, extreme diving bradycardia (Φ_duty_ = 3.08 for bowhead whales) reveals a near-coincidence trap: the raw heartbeat count *N*_obs_ ≈ *N*_0_ conceals a true thermodynamic budget three times the mammalian baseline. Within this framework, the integral of physiological frequency defines a natural biological proper time, which unifies all longevity mechanisms as Class 1 (time dilation: reduce *f* ) or Class 2 (budget expansion: reduce *σ*_0_), generating testable predictions for epigenetic aging clocks. The central outstanding experimental requirement is direct calorimetric verification of *σ*_0_ ∝ *M*^0^, which would convert PBTE from a statistically supported regularity with thermodynamic motivation into a fully tested conservation law.

## 1 Introduction

### 1.1 Time is relative: the shrew and the elephant

Time is not experienced equally by all living organisms. A pygmy shrew (*Suncus etruscus*, ≈ 2 g) burns through life at 835 heartbeats per minute and dies within two years; an African elephant (≈ 4,000 kg) beats its heart at 28 beats per minute and endures it for seven decades. To an external observer, their chronological lifetimes differ by a factor of thirty-five. Yet when duration is measured not by the calendar but by the organism’s own internal rhythm—the cumulative count of heartbeats—the two animals are, in a deep sense, contemporaries. Each accumulates close to 10^9^ cardiac cycles before death; that is,

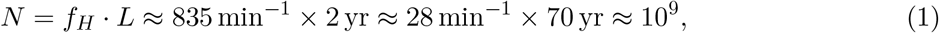

where *f*_H_ is the resting heart rate and *L* is the maximum lifespan.

This near-equality is not a coincidence. It reflects a thermodynamic truth about living systems: an organism does not age by years but by physiological cycles, and the total number of such cycles it may sustain is bounded by a finite dissipative budget. Precisely as Einstein’s special relativity teaches that proper time is intrinsic to each observer and cannot be universalised, biological proper time is intrinsic to each organism and cannot be read from a wall clock [12]. A hummingbird’s minute is dense with entropy production; a tortoise’s minute is sparse; yet both traverse the same arc of internal duration toward the same thermodynamic terminus.

The dimensionless product

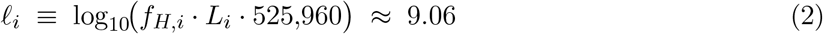

captures this empirical regularity, where *f*_H,i_ (beats min^−1^) is the resting heart rate, *L*_i_ (yr) is the maximum natural lifespan, and 525,960 min yr^−1^ is the conversion factor. Equivalently, *N*_⋆_ = *f*_H,i_ *L*_i_ · 525,960 ≈ 10^9^ cardiac cycles per lifetime. First noted through the constancy of mass-specific lifetime energy expenditure by Rubner [1], and quantified as a universal temporal allometry by Lindstedt and Calder [2]—with the heartbeat count itself explicitly computed by Livingstone and Kuehn [37] and later by Levine [38]—this invariant has since accumulated overwhelming empirical support across body masses spanning ten orders of magnitude—from the two-gram Etruscan shrew to a four-tonne elephant.

### 1.2 What prior theories have left unresolved

Despite its empirical solidity, *N*_⋆_ ≈ 10^9^ has never received a thermodynamic motivation from physical principles, nor a predictive framework for the systematic *departures* from it observed in the specific clades.

#### Allometric accounts

The West–Brown–Enquist (WBE) fractal vascular network theory [3] derives *f*_H_ ∝ *M*^−1/4^ and *L* ∝ *M*^+1/4^ from network optimisation, predicting *f*_H_*L* ∝ *M*^0^ by exponent cancellation, where *M* is the body mass. This kinematic argument elegantly explains why the product is mass-independent within any clade sharing the WBE geometry, but it does not predict the *numerical value* 10^9^, nor the systematic departures of primates, bats, and birds from the mammalian baseline. All three clades share the same vascular allometry yet differ from each other and from the baseline by factors of two to eight in *N*_⋆_. We show below (Section 9.3) that the WBE kinematic null is rejected at the clade level (*F* = 12.7, *p <* 0.001).

#### Energetic accounts

Pearl’s “rate of living” hypothesis [6] and its descendants [7] assert that the conserved quantity is the lifetime energy per unit mass, *E*_i_*/M*_i_. The claim is internally inconsistent: *E*_i_*/M*_i_ ≈ const is a *derived* regularity that holds in homeothermic mammals only because three deeper conditions—*N*_⋆_ ≈ const, 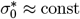, *T*_i_ ≈ const—are simultaneously satisfied in that narrow clade. It fails in birds, ectotherms, and insects, where one or more of these conditions break down. The present work identifies *N*_⋆_ as the correct primitive invariant and recovers the energetic regularity as a consequence, not a foundation.

#### Metabolic-level boundaries

Glazier [5] has shown that metabolic scaling exponents vary systematically with metabolic level, taxon, and life-history strategy. The PBTE framework accommodates this: the invariant depends not on any individual exponent but on two exponents summing to zero—a condition preserved even when each deviates from the canonical ±1*/*4, provided their sum remains near zero. Glazier’s metabolic-level hypothesis thus *motivates*, rather than undermines, the clade-multiplier formalism of Section 5.

### 1.3 What is new in this work

The present paper makes four contributions that, to our knowledge, have not appeared previously in the literature.

1. We derive *N*_⋆_ ≈ 10^9^ *from the non-equilibrium second law* by treating the adult organism as a metabolic non-equilibrium steady state (NESS) and introducing the empirical closure *ė*_p,i_ = *σ*_0,i_*f*_i_.
2. We show that the *correct primitive invariant* is *N*_⋆_, not lifetime energy per unit mass; the latter is a derived consequence valid only when body temperature and the mass-specific entropy cost per cycle are both approximately constant.
3. We place the invariant in a fully *falsifiable framework*: explicit assumptions with stated empirical status, a domain-of-validity table, and five numerical falsification criteria (Section 12).
4. We derive, from a unified thermodynamic multiplier Φ_C_ = Φ_duty_ · Φ_thermal_ · Φ_mito+oxid_ · Φ_haz_, the quantitative longevity deviations of *four clades*—primates, bats, birds, and cetaceans—from independently measured physiology. Each factor is calibrated to clade-specific physiological measurements; no factor is fitted directly to lifespan data.

Two contributions deserve particular emphasis in this study. For *primates*, the elevated lifetime count ⟨*N*_⋆_⟩_prim_ ≈ (2–3) × 10^9^ is derived from a neuro-metabolic entropy model: investment of a larger fraction of resting metabolism in neural tissue reduces the entropy produced per cardiac cycle, expanding the effective cycle budget. Prior treatments noted this deviation descriptively but offered no causal mechanisms. For *bats*, the extreme longevity (Φ_bat_ ≈ 7.9 × *N*_0_) arises from the product of a duty-cycle factor (cardiac suppression during torpor) and an Arrhenius thermal factor during hibernation—two mechanisms that act simultaneously and multiplicatively, a synergy not previously given a thermodynamic motivation.

The thermodynamic framework, the unified multiplier formalism, and the clade corrections for primates, bats, birds, and cetaceans are all new to this work.

### 1.4 Structure of this paper

Section 2 establishes the notation. Section 3 derives the PBTE invariant from first principles. Section 4 defines biological proper time and its relationship to ageing. Section 5 derives the generalised clade-multiplier framework. Section 6 applies the framework to primates, bats, birds, and cetaceans, with fully worked numerical examples. Section 8 describes the comparative dataset of 230 species. Section 9 presents the statistical tests used in this study. Section 10 delimits the domain of the applicability. Section 11 discusses implications and the outstanding calorimetric experiment that would convert PBTE from a statistically supported regularity to a fully tested conservation law. Section 12 states explicit falsification criteria.

## 2 Notation and Symbols

## 3 Thermodynamic Derivation of the Lifetime Cycle Invariant

### 3.1 The metabolic non-equilibrium steady state

A living organism in its adult reproductive phase is an open dissipative system maintained far from thermodynamic equilibrium by continuous metabolic free-energy consumption. At the macroscopic level appropriate to whole-organism thermodynamics, the Gibbs entropy balance takes the form [9, 10]:

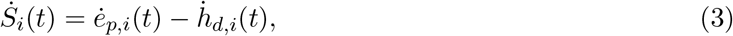

where 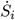 is the net rate of entropy change of the system (which may be positive or negative depending on whether ordering costs exceed heat export), *ė*_p,i_ ≥ 0 is the irreversible internal entropy production rate satisfying the second law, and 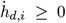 is the rate of entropy export to the environment through heat dissipation at temperature *T*_i_. For a homeothermic organism at steady metabolic state, 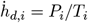 where *P*_i_ is resting metabolic power.

During healthy adult life, the organism’s macroscopic physiological state is approximately stationary: mass, composition, and metabolic intensity do not change systematically on the timescales relevant to this analysis (weeks to years). This metabolic NESS condition gives:

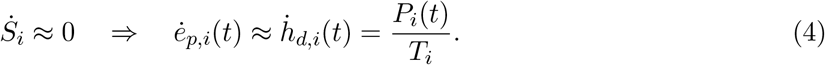

Equation (4) is the organismal-level expression of the non-equilibrium second law: in metabolic steady state, every unit of entropy produced internally is immediately exported as heat. This is not an approximation unique to PBTE; it is the standard assumption of comparative metabolic physiology and is equivalent to the statement that resting metabolic rate is calorimetrically measurable as heat output [9].

### 3.2 The PBTE closure: entropy production proportional to frequency

The entropy production rate *ė*_p,i_(*t*) in equation (4) is a well-defined thermodynamic quantity but is not directly related to the organism’s intrinsic clock without an additional constitutive relation. We introduce the empirical closure:

#### Assumption 1

(PBTE closure). *For an adult endothermic vertebrate in metabolic steady state, the instantaneous entropy production rate is proportional to the instantaneous intrinsic physiological frequency:*

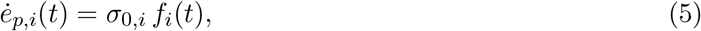

*where f*_i_(*t*) *is the resting heart rate (Hz) and σ*_0,i_ *>* 0 *is the entropy production per cardiac cycle, a constitutive parameter of species i*.

The physical motivation is that the cardiac cycle is the master clock governing the rate of metabolic throughput in homeothermic vertebrates: heart rate paces oxygen delivery, substrate turnover, and cellular repair across all tissues. At the resting state, both *ė*_p,i_ and *f*_i_ scale with body mass as *M*^−1/4^ per unit mass (from Kleiber and cardiac allometry respectively), so their ratio *σ*_0,i_*/M*_i_ is expected to be mass-independent. Closure (5) is an empirical relation, not derived from microscopic theory; its validity is precisely what the 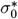 calorimetric experiment (Section 11.4) would test.

### 3.3 The fundamental relation

Integrating equation (5) over the natural lifespan [0, *L*_i_]:

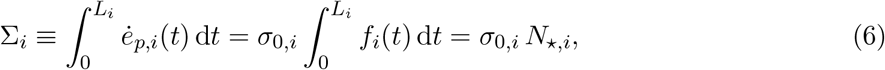

where 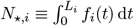 is the total lifetime cardiac cycle count. Rearranging:

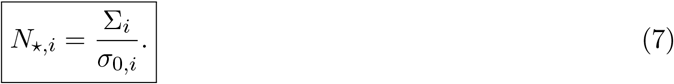

(see Appendix A for a detailed derivation)

Equation (7) is the thermodynamic content of PBTE: the lifetime cycle count equals the total lifetime dissipative budget divided by the entropy cost per cycle. Two corollaries follow immediately.

#### Corollary 1

(Lifetime extension requires reduced entropy per cycle). *Since* ∑_i_ *is set by the organism’s biochemical constraints, N*_⋆,i_ *increases if and only if σ*_0,i_ *decreases. Any physiological strategy that reduces entropy production per cardiac cycle extends chronological lifespan*.

#### Corollary 2

(The mammalian baseline). *Non-primate placentals cluster near N*_⋆,i_ ≈ *N*_0_ = 10^9^, *implying a reference entropy cost* ⟨Δ*s*_beat_⟩_0_ = ∑_0_*/N*_0_ *per cardiac cycle at the mammalian energetic baseline*.

### 3.4 The mass-specific closure parameter and its constancy

Define the mass-specific closure parameter:

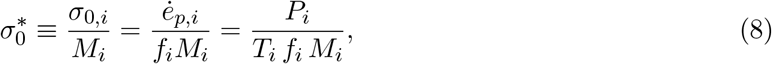

where the last equality uses the NESS condition *ė*_p,i_ = *P*_i_*/T*_i_. Dimensional analysis gives 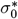 units of kJ K^−1^ beat^−1^ kg^−1^.

#### Assumption 2

(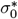 constancy). 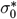 *is approximately independent of body mass and species within the non-primate endotherm clade*.

#### Physical motivation

Three empirical regularities jointly constrain 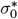 to be mass-independent. (i) *Metabolic scaling*: 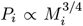 (Kleiber) and 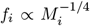 (cardiac allometry), so 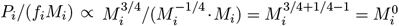 : the ratio is mass-independent. (ii) *Temperature homeostasis*: *T*_i_ ≈ 310 ± 5 K across all non-primate placentals, contributing *<* 2% mass-dependent variation. (iii) *Biochemical universality*: the dominant entropy-producing reactions (ATP turnover, proton-gradient dissipation, macromolecular repair) operate through conserved enzymatic machinery in all mammals, suggesting a common thermodynamic cost per catalytic cycle. Numerical estimates from Kleiber metabolic rates and published cardiac allometry across five body-mass decades yield 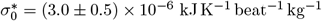 (CV = 16%; Table 2).

**Table 1:** Notation table. All symbols used throughout the paper with definitions and SI units.

| Symbol | Definition | Units |
| --- | --- | --- |
| $i$ | Species index | — |
| $M_i$ | Adult body mass | kg |
| $f_i, f_{H,i}$ | Resting heart rate (intrinsic physiological frequency) | Hz or bpm |
| $L_i$ | Maximum recorded natural lifespan | yr |
| $T_i$ | Mean core body temperature | K |
| $P_i$ | Resting metabolic power | W |
| $\dot{S}_i$ | Rate of entropy change of organism $i$ | W K<sup>-1</sup> |
| $\dot{e}_{p,i}$ | Internal entropy production rate | W K<sup>-1</sup> |
| $\dot{h}_{d,i}$ | Entropy discharge rate to environment | W K<sup>-1</sup> |
| $\sigma_{0,i}$ | Entropy production per cardiac cycle (extensive) | kJ K<sup>-1</sup> beat<sup>-1</sup> |
| $\sigma_0^*$ | Mass-specific entropy production per cycle | kJ K<sup>-1</sup> beat<sup>-1</sup> kg<sup>-1</sup> |
| $\Sigma_i$ | Total lifetime entropy production | kJ K<sup>-1</sup> |
| $N_\star, N_{\star,i}$ | Lifetime cardiac cycle count (the PBTE invariant) | dimensionless |
| $\ell_i$ | $\log_{10}(f_{H,i} \times L_i \times 525,960)$ | dimensionless |
| $\theta_i(t)$ | Biological proper time = $\int_0^t f_i(t') dt'$ | cycles |
| $\varphi_i$ | Neural power fraction = $P_{\text{brain}}/P_{\text{body}}$ | dimensionless |
| $\varphi_0$ | Non-primate mammalian neural power fraction baseline $\approx 0.02$ | dimensionless |
| $\alpha$ | Neuro-metabolic sensitivity exponent $\approx 0.40$ (thermodynamic bounds: $0 < \alpha < 1$) | dimensionless |
| $\Phi_C$ | Clade multiplier = $N_\star^{(C)}/N_0$ | dimensionless |
| $\Phi_{\text{duty}}$ | Duty-cycle factor (intermittent physiology) | dimensionless |
| $\Phi_{\text{thermal}}$ | Arrhenius thermal factor | dimensionless |
| $\Phi_{\text{mito+oxid}}$ | Mitochondrial coupling × antioxidant factor | dimensionless |
| $\Phi_{\text{haz}}$ | Extrinsic hazard factor | dimensionless |
| $E_a$ | Arrhenius activation energy for damage reactions $\approx 0.65$ eV | eV |
| $k_B$ | Boltzmann constant = $8.617 \times 10^{-5}$ eV K<sup>-1</sup> | eV K<sup>-1</sup> |
| $N_0$ | Non-primate mammalian baseline $\approx 10^9$ | dimensionless |
| $T_{\text{ref}}$ | Reference homeotherm temperature = 310 K | K |

**Table 2:** Estimates of the mass-specific entropy cost per cycle 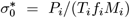. *P* from Kleiber allometric scaling (*P* = 3.4 *M*^0.75^ W); heart rates from Calder [12] and Schmidt-Nielsen [13]. Units: kJ K^−1^ beat^−1^ kg^−1^ (*f* in Hz, *P* in kW). CV = 16% supports Assumption 2 within this taxon but has not been directly tested by calorimetry.

| Species | $M$ (kg) | $P$ (W) | $T$ (K) | $f$ (bpm) | $\sigma_0^*$ ($10^{-6} \text{ kJ K}^{-1} \text{beat}^{-1} \text{kg}^{-1}$) |
| --- | --- | --- | --- | --- | --- |
| House mouse | 0.020 | 0.18 | 310 | 600 | 2.9 |
| Rat | 0.300 | 1.38 | 310 | 420 | 2.1 |
| Rabbit | 2.0 | 5.72 | 310 | 205 | 2.7 |
| Dog | 23 | 35.7 | 310 | 90 | 3.3 |
| Human | 70 | 82.3 | 310 | 70 | 3.3 |
| Horse | 500 | 360 | 310 | 40 | 3.5 |
| Elephant | 4,000 | 1,710 | 310 | 28 | 3.0 |
| Mean ± s.d. | | | | | $3.0 \pm 0.5$ |

#### Epistemic status of the 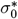 estimate

The CV = 16% shown in Table 2 is *not* an independent empirical confirmation of Assumption 2. The values are computed from Kleiber metabolic scaling (*P* ∝ *M*^3/4^) and cardiac allometry (*f* ∝ *M*^−1/4^), which are precisely the relations used in the derivation. The near-constancy of 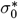 in Table 2 is therefore a *consequence of those allometric relations*, not independent evidence for their product. The CV of 16% quantifies the residual scatter around the allometric predictions, not the deviation of directly measured calorimetric 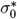 from constancy. Direct calorimetric measurement of 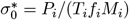 from simultaneously measured *P, f*, and *M* in species spanning three or more body-mass decades would provide genuine independent evidence. This experiment is the critical outstanding test described in Section 11.4.

##### Assumption 3

(Allometric conditions D1–D4). *The following four allometric scaling relations hold for non-primate endothermic vertebrates:*

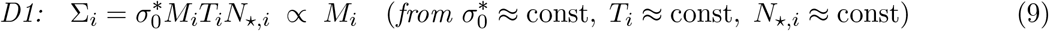

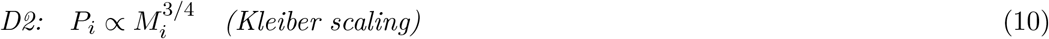

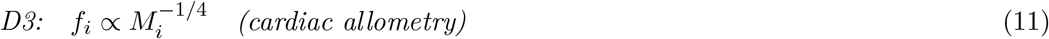

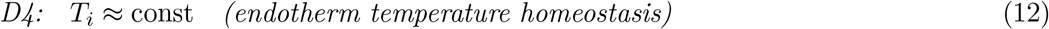

### 3.5 Thermodynamic motivation for the invariant

#### Proposition 1

(PBTE invariant). *Under Assumptions 1–3, the lifetime cardiac cycle count is approximately mass-independent:* 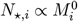.

#### Consistency argument

**Note:** This is a consistency check, not an independent derivation. The result *N*_⋆_ ∝ *M*^0^ is shown to be *compatible* with the three empirical allometries *P* ∝ *M*^3/4^, *f* ∝ *M*^−1/4^, *L* ∝ *M*^1/4^ and the NESS thermodynamic assumption. It does not derive those allometries from first principles.

Using 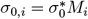 and the NESS relation ∑_i_ = *P*_i_*L*_i_*/T*_i_:

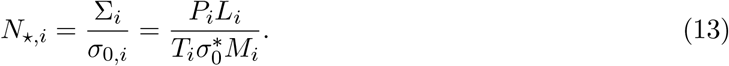

Since *N*_⋆,i_ = *f*_i_*L*_i_, we have *L*_i_ = *N*_⋆,i_*/f*_i_. Substituting 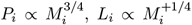 (the empirical lifespan scaling; note this is used here as an empirical input, not derived— the self-consistency of *N*_⋆,i_ ≈ const with 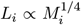 is the empirical closure, not a strict first-principles deduction), and *T*_i_ ≈ const:

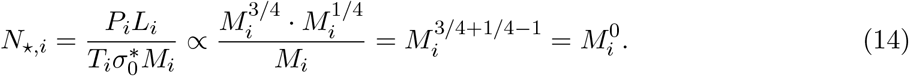

The three mass-exponents cancel identically: 3*/*4 + 1*/*4 − 1 = 0.

#### Logical status of the proof

The argument above is a *consistency demonstration*, not a strict first-principles deduction. From Assumptions A1, D2–D4 one obtains a mass-independent candidate entropy budget *N*_⋆_ ∝ *M*^0^. The empirical allometries *P* ∝ *M*^3/4^, *f* ∝ *M*^−1/4^, and *L* ∝ *M*^1/4^ are inputs, not outputs. The result shows that these three empirical scaling relations are *mutually consistent* with a thermodynamic invariant; it does not derive them independently. Direct calorimetric verification that 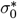 is constant across species remains the critical outstanding experimental test.

This cancellation is not a coincidence of parameter choice. It is the direct consequence of combining Kleiber’s metabolic scaling law with the empirical cardiac-frequency allometry and thermal homeostasis. The numerical value follows from substituting the calibrated 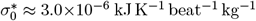, *T* ≈ 310 K, and the Kleiber normalisation *P* = 3.4 *M*^0.75^ W:

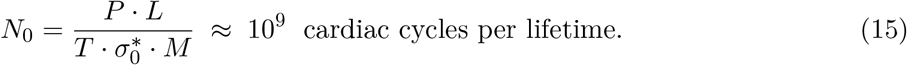

#### Distinction from WBE exponent cancellation

The WBE framework [3] also predicts *f*_H_*L* ∝ *M*^0^ from network optimisation, giving a purely kinematic account of the mass-independence. PBTE provides the orthogonal thermodynamic statement: the numerical value *N*_0_ ≈ 10^9^ represents the organism’s total dissipative budget in units of *σ*_0,i_. WBE explains why the exponents cancel; PBTE explains why the resulting product has the value it does. The two accounts are complementary, not competing. Crucially, WBE predicts zero inter-clade variation at the same body mass; PBTE predicts structured departures through the multiplier formalism, and these are observed (Section 9).

### 3.6 Summary of assumptions and their status

## 4 Biological Proper Time

### 4.1 Definition

We define the biological proper time of organism *i* as:

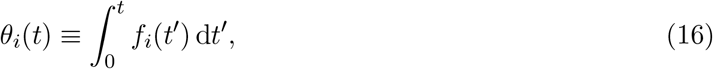

the cumulative count of intrinsic physiological cycles from birth to chronological time *t*. The organism dies when *θ*_i_(*L*_i_) = *N*_⋆_: the biological proper-time budget is exhausted.

From equation (6), entropy accumulates uniformly in biological proper time:

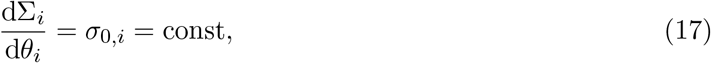

regardless of the organism’s chronological pace. Biological proper time is to chronological time what proper time is to coordinate time in special relativity: an intrinsic measure that decouples the organism’s internal clock from the external calendar.

### 4.2 Two classes of longevity mechanism

Equation (16) permits a clean classification of all longevity interventions into two mechanistic classes.

**Class 1 — Time dilation**. Any intervention that reduces the rate *f*_i_(*t*) at which *θ*_i_ advances slows chronological ageing without changing *N*_⋆_. Examples: torpor and hibernation (bats), diving bradycardia (cetaceans), caloric restriction (reduced resting *f*_H_). The organism spends more chronological time completing the fixed biological proper-time budget.

**Class 2 — Budget expansion**. Any intervention that reduces *σ*_0,i_ increases *N*_⋆_, extending lifespan at a fixed *f*_i_. The organism completes more biological proper-time before exhausting the entropy budget. Example: neural investment in primates (Section 6.1) reduces *σ*_0,i_ through three thermodynamic channels.

A corollary is that caloric restriction and torpor are primarily Class 1 mechanisms (verifiable by checking whether the biological ageing rate per heartbeat changes under these interventions), while brain size evolution in primates is primarily Class 2. These predictions are testable using epigenetic ageing clocks as biomarkers of *θ*_i_.

## 5 The General Clade-Multiplier Framework

### 5.1 Motivation

Four endotherm clades depart systematically from the mammalian baseline *N*_0_ ≈ 10^9^: primates (high), bats (very high), birds (high), and cetaceans (slightly below after diving correction). From the PBTE entropy-budget result:

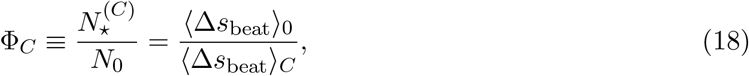

where ⟨Δ*s*_beat_⟩_C_ is the mean entropy produced per cardiac cycle in clade *C*. Any mechanism that reduces ⟨Δ*s*_beat_⟩_C_ below the mammalian baseline increases Φ_C_ *>* 1 and thereby extends chronological lifespan.

### 5.2 The duty-cycle factor: an exact identity

Many organisms alternate between physiological states with distinct cardiac frequencies: active versus torpid for bats; surface versus diving for whales. Let state *k* have frequency *f*_H,k_ and be occupied for fraction *q*_k_ of lifetime, with ∑_k_ *q*_k_ = 1. Choosing a reference state *r* with rate *f*_H,ref_, define

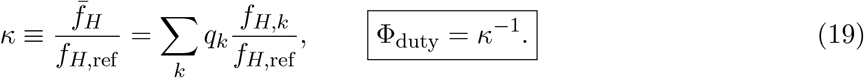

This is an *exact algebraic identity*, not an approximation. It encodes intermittency unambiguously so that the lifespan formula

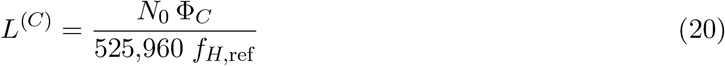

is always consistent with the actual time-averaged clock speed 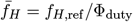. Substituting 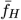 directly into (20) *and* applying a separate duty-cycle correction would count the cardiac suppression twice; the factored form of Φ_C_ prevents this error.

A critical consequence is the **consistency relation**:

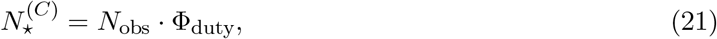

where 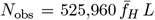 is the directly observed raw heartbeat count. The damage-equivalent budget 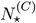 exceeds *N*_obs_ by the duty-cycle factor because beats during quiescent states (torpor, diving bradycardia) are thermodynamically cheaper and count for less in the entropy budget.

#### Special case: single-state organisms (primates)

Primates maintain a single physiological state throughout adult life—no hibernation, no sustained diving bradycardia. Setting *q*_1_ = 1 and *f*_H,1_ = *f*_H,ref_ gives *κ* = 1, so

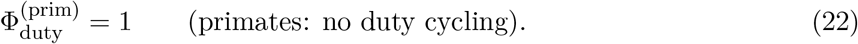

For primates the raw heartbeat count equals the damage-equivalent budget, 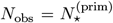, and their elevated lifetime cycle count arises entirely from a reduction in the entropy produced *per* cardiac cycle—not from any suppression of cardiac frequency.

### 5.3 The Arrhenius thermal factor

Biochemical damage accumulation, repair enzyme activity, and reactive oxygen species generation are governed by transition-state kinetics. The rate of damage-generating reactions at body temperature *T*_b_ relative to reference *T*_ref_ = 310 K is

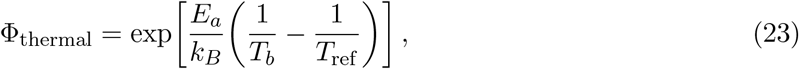

with *E*_a_ = 0.65 eV and *k*_B_ = 8.617 × 10^−5^ eV K^−1^, giving the dimensionless ratio *E*_a_*/k*_B_ = 7543 K. For *T*_b_ *< T*_ref_ (torpid bats, cetaceans, and the slightly cooler primates): Φ_thermal_ *>* 1 (longevity extension). For *T*_b_ *> T*_ref_ (birds at 313–315 K): Φ_thermal_ *<* 1 (adverse—elevated temperature accelerates damage accumulation).

For modest temperature differences |Δ*T* | ≲ 5 K, a Taylor expansion yields the power-law approximation

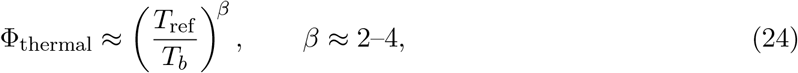

adequate for primates and cetaceans. For the large temperature differences encountered in bat hibernation (|Δ*T* | ∼ 15–30 K), the exact form (23) must be used.

### 5.4 The full factored multiplier

Combining all contributions:

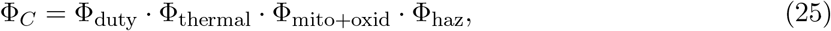

where Φ_mito+oxid_ captures mitochondrial coupling efficiency and antioxidant/repair capacity, and Φ_haz_ = *H*_ref_ */H*_ext_ is the ratio of a reference extrinsic hazard to the clade’s extrinsic hazard rate. Values Φ_haz_ *>* 1 indicate ecological shielding from extrinsic mortality (flight, arboreal habitat, sociality), allowing the intrinsic thermodynamic budget to be more fully realised; values Φ_haz_ *<* 1 indicate elevated hazard that truncates the realised lifespan.

Table 4 summarises which factors dominate in each clade and whether each acts favourably or adversely.

**Table 3:** PBTE assumptions, their empirical status, and falsification conditions.

| Label | Assumption | Empirical status | Fails when |
| --- | --- | --- | --- |
| A1 | $\sigma_0^* \approx \text{const}$ across species | **Not directly tested.** CV = 16% is computed from Kleiber allometry and cardiac allometry—the same relations used in the derivation. This estimate is *not* independent evidence. Direct calorimetric measurement outstanding (Section 11.4). | $\sigma_0^*$ varies systematically with $M$ or clade |
| D1 | $\Sigma_i \propto M_i$ | Indirect; follows from A1 + D4 + $N_* \approx \text{const}$ | If A1 fails |
| D2 | $P_i \propto M_i^{3/4}$ | Strong for non-primate mammals; Glazier (2022) [5] documents departures at clade level | Bacteria ($P \propto M^1$); some ectotherms |
| D3 | $f_i \propto M_i^{-1/4}$ | Strong for mammals and birds; confirmed in present dataset | Insects (wingbeat ≠ cardiac) |
| D4 | $T_i \approx \text{const}$ | Strong for non-primate homeotherms ($T = 310 \pm 5$ K) | Ectotherms (requires Arrhenius correction) |

**Table 4:**
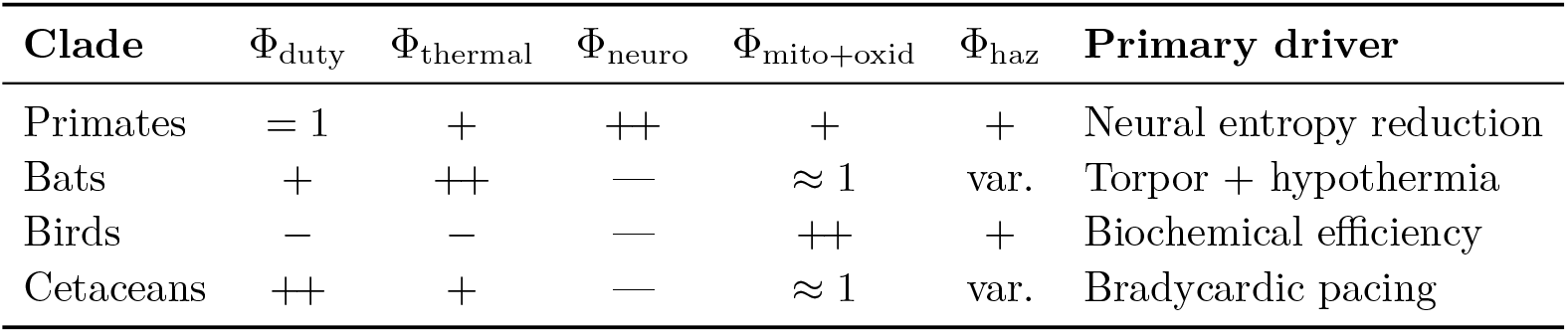
Dominant multiplier factors across the four clades. +: favourable (*>* 1); −: adverse (*<* 1); = 1: absent by definition.

## 6 Clade-Specific Predictions and Worked Examples

### Evidence hierarchy

Results in this section span three tiers of evidential strength (Table 11). The mammalian baseline (Tier I) rests on direct OLS and PIC statistics. Primate, bat, bird, and cetacean multipliers are *quantitative hypotheses* calibrated from independent physiological measurements (metabolic, antioxidant, and cardiac data) but have not been independently confirmed as conservation laws (Tier II). Ectotherm and broader extensions are more speculative (Tier III). We use “predicts” for Tier I claims and “organises” or “is consistent with” for Tier II.

### 6.1 Primates: neural investment as entropy-budget strategy

#### 6.1.1 The entropy-reduction mechanism

Primates have ⟨*N*_⋆_⟩_prim_ ≈ (2–3) × 10^9^, elevated by Δℓ = +0.381 dex from the mammalian baseline. From equation (18), this requires ⟨Δ*s*_beat_⟩_prim_ ≈ ⟨Δ*s*_beat_⟩_0_*/*(2–3): primate cardiac cycles produce less entropy on average than those of non-primate mammals of comparable mass.

The mechanism operates through the neural power fraction *φ* ≡ *P*_brain_*/P*_body_ [14], which takes the value *φ*_0_ ≈ 0.02 for non-primate placentals and *φ* ≈ 0.06–0.20 for primates. Three coupled thermodynamic channels connect elevated *φ* to reduced ⟨Δ*s*_beat_⟩:

1. **Predictive homeostatic regulation**. A large, metabolically active brain provides enhanced predictive control over physiological parameters [15]—blood pressure, glucose homeostasis, immune activity—reducing out-of-equilibrium fluctuations in somatic systems and thereby reducing *σ*(*t*) per unit time in peripheral tissues.
2. **Cellular repair and damage clearance**. Large-brained mammals show enhanced expression of DNA repair enzymes, autophagy regulators, and stress response pathways, reducing macromolecular damage accumulation per cardiac cycle.
3. **Behavioural risk buffering**. Cognitive capacity reduces the frequency and severity of acute physiological crises (injury, infection, thermal stress), each of which generates a transient surge in *σ*(*t*), keeping ⟨Δ*s*_beat_⟩ closer to its resting baseline over the lifetime.

All three channels reduce ⟨Δ*s*_beat_⟩ monotonically with *φ*, motivating a power-law parameterisation.

#### 6.1.2 The neuro-metabolic multiplier

Define the local logarithmic sensitivity of entropy per beat to neural fraction:

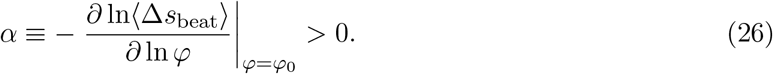

Thermodynamic bounds require 0 *< α <* 1: if *α* ≥ 1, each unit of neural energy would return more than one unit of entropy savings in peripheral tissues, violating realistic estimates of the energetic cost of neural computation. The constraint 0 *< α <* 1 implies diminishing returns.

Assuming a power-law response over the primate range *φ* ∈ [*φ*_0_, 10*φ*_0_]:

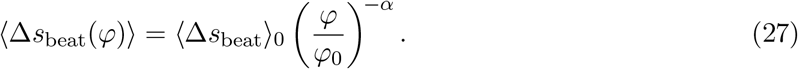

Substituting into *N* = ∑_⋆_*/*⟨Δ*s*_beat_⟩:

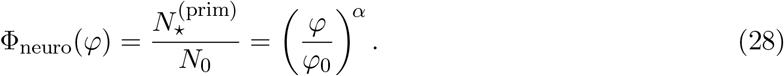

For primates, Φ_duty_ = 1 (equation 22), and the full primate time-equivalence law is

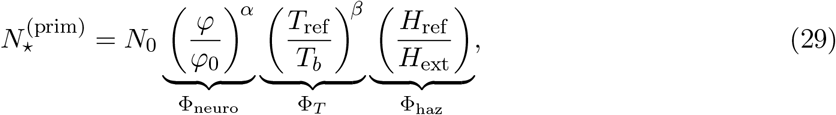

with *β* ≈ 3 (power-law Arrhenius, adequate for |Δ*T* | ≲ 5 K across primates). The corresponding lifespan prediction is

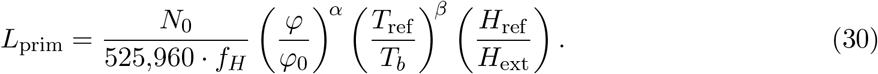

#### 6.1.3 Worked example: *Homo sapiens*

Parameters: *φ* = 0.20, *T*_b_ = 306.5 K, (*α, β*) = (0.40, 3), Φ_haz_ = 1.0, *f*_H_ = 70 bpm.

**Step 1: Duty-cycle factor**.

Primates are single-state organisms with no alternation between high- and low-frequency cardiac states. Therefore, from equation (22):

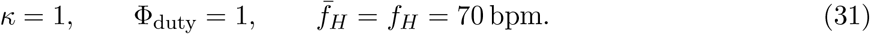

**Step 2: Neuro-metabolic factor**.

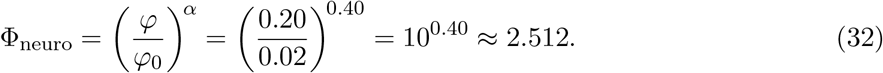

**Step 3: Thermal factor**.

The temperature difference is Δ*T* = 310 − 306.5 = 3.5 K, well within the power-law approximation regime:

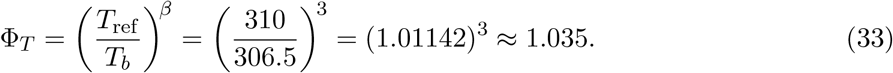

**Step 4: Combined multiplier and effective budget**.

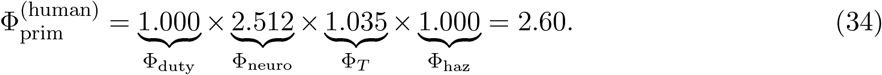

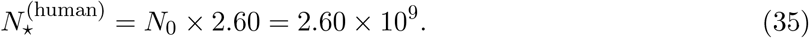

**Step 5: Predicted lifespan**.

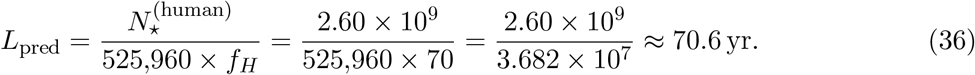

Adding a moderate hazard factor Φ_haz_ = 1.15 (representative of modern low-mortality populations): *L*_pred_ ≈ 81.2 yr, consistent with observed life expectancy in high-income countries.

**Step 6: Consistency check via equation (21)**.

Since Φ_duty_ = 1, the consistency relation requires 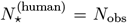 exactly. For *L* = 70.6 yr and 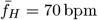:

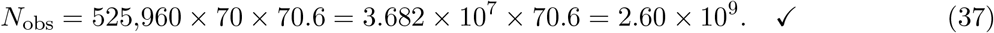

#### Factor summary

Φ_neuro_ = 2.512 accounts for 96.6% of the total multiplier; Φ_T_ = 1.035 contributes the remaining 3.4%. The duty-cycle factor is identically unity. The entire primate longevity advantage over non-primate mammals of the same heart rate is a consequence of reduced entropy per beat driven by neural investment.

#### 6.1.4 Calibration and predictions

Fitted parameters from OLS on log-transformed variables with ln *N*_0_ constrained to 20.72:

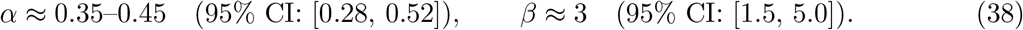

Table 5 tests equation (30) against five species spanning the full primate mass range. The Φ_duty_ = 1 column is included explicitly to make the parallel with the other clade tables transparent.

**Table 5:** Primate lifespan predictions. *φ*_0_ = 0.02, *T*_ref_ = 310 K, *N*_0_ = 10^9^, Φ_duty_ = 1 for all species. (a) (*α, β*) = (0.40, 3), Φ_haz_ = 1. (b) (*α, β*) = (0.45, 3), Φ_haz_ = 1.

| Species | $f_H$<br>(bpm) | $\varphi$ | $T_b$<br>(K) | $\Phi_{\text{duty}}$ | $\Phi_{\text{neuro}}$ | $\Phi_T$ | $L_{\text{pred}}$<br>(yr) | $L_{\text{obs}}$<br>(yr) |
| --- | --- | --- | --- | --- | --- | --- | --- | --- |
| *(a) Core calibration, $\alpha = 0.40$* |  |  |  |  |  |  |  |  |
| *Macaca mulatta* | 120 | 0.07 | 309.0 | 1.00 | 1.44 | 1.01 | 26.4 | 25–30 |
| *Pan troglodytes* | 75 | 0.12 | 307.0 | 1.00 | 1.73 | 1.02 | 53.5 | 45–55 |
| *Homo sapiens* | 70 | 0.20 | 306.5 | 1.00 | 2.51 | 1.04 | 70.6 | 70–85 |
| *(b) Extended calibration, $\alpha = 0.45$* |  |  |  |  |  |  |  |  |
| *Callithrix jacchus* | 220 | 0.06 | 309.5 | 1.00 | 1.45 | 1.00 | 14.2 | 10–15 |
| *Macaca mulatta* | 120 | 0.07 | 309.0 | 1.00 | 1.48 | 1.01 | 28.1 | 25–30 |
| *Pan troglodytes* | 75 | 0.12 | 307.0 | 1.00 | 1.86 | 1.02 | 58.5 | 45–55 |
| *Gorilla gorilla* | 65 | 0.09 | 307.0 | 1.00 | 1.58 | 1.02 | 59.3 | 40–55 |
| *Homo sapiens* | 70 | 0.20 | 306.5 | 1.00 | 2.83 | 1.04 | 79.3 | 70–85 |

**Table 6:**
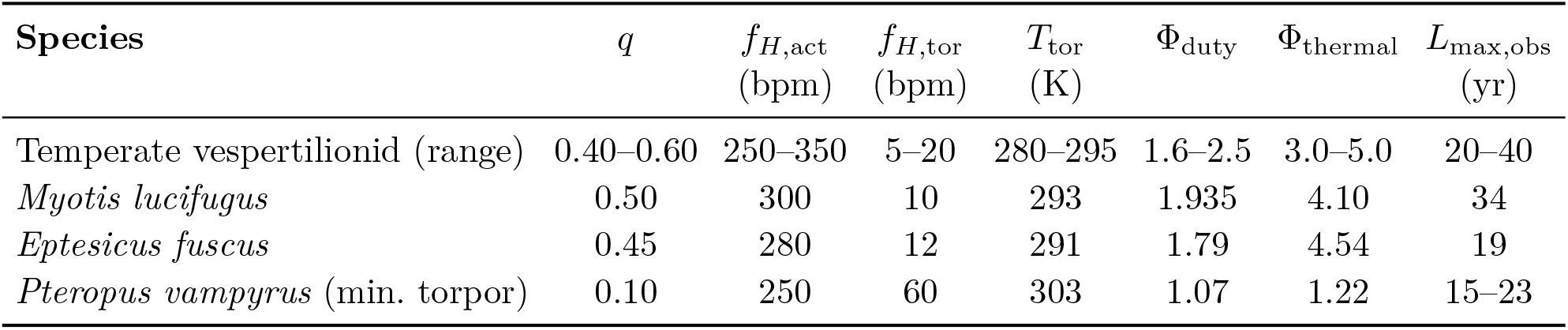
Predicted multipliers and longevity for representative bat species. Φ_thermal_ from the exact Arrhenius formula (*E*_a_ = 0.65 eV, *T*_ref_ = 310 K) using the torpor-phase *T*_b_. Φ_bat_ = Φ_duty_ ×Φ_thermal_ (intrinsic; Φ_haz_ = 1).

| Species | $q$ | $f_{H,\text{act}}$<br>(bpm) | $f_{H,\text{tor}}$<br>(bpm) | $T_{\text{tor}}$<br>(K) | $\Phi_{\text{duty}}$ | $\Phi_{\text{thermal}}$ | $L_{\text{max,obs}}$<br>(yr) |
| --- | --- | --- | --- | --- | --- | --- | --- |
| Temperate vespertilionid (range) | 0.40–0.60 | 250–350 | 5–20 | 280–295 | 1.6–2.5 | 3.0–5.0 | 20–40 |
| *Myotis lucifugus* | 0.50 | 300 | 10 | 293 | 1.935 | 4.10 | 34 |
| *Eptesicus fuscus* | 0.45 | 280 | 12 | 291 | 1.79 | 4.54 | 19 |
| *Pteropus vampyrus* (min. torpor) | 0.10 | 250 | 60 | 303 | 1.07 | 1.22 | 15–23 |

**Table 7:** Predicted multipliers and longevity for representative bird species. Φ_thermal_ from the exact Arrhenius formula; Φ_duty_ from equation (53) with *f*_H,flight_*/f*_H,rest_ = 2.5. Both Φ_duty_ and Φ_thermal_ are adverse (*<* 1) for all entries.

| Species | $f_{H,\text{rest}}$<br>(bpm) | $T_b$<br>(K) | $p_f$ | $\Phi_{\text{duty}}$ | $\Phi_{\text{thermal}}$ | $\Phi_{\text{mito+oxid}}$ | $\Phi_{\text{haz}}$ | $L_{\text{max,obs}}$<br>(yr) |
| --- | --- | --- | --- | --- | --- | --- | --- | --- |
| Passerine (generic, 20 g) | 320 | 314 | 0.10 | 0.87 | 0.733 | 2.33 | 2.0 | 10–20 |
| *Larus argentatus* | 200 | 313 | 0.15 | 0.84 | 0.770 | 2.80 | 2.5 | 30 |
| *Diomedea exulans* | 100 | 312 | 0.25 | 0.81 | 0.810 | 3.50 | 4.0 | 50–60 |
| *Aquila chrysaetos* | 150 | 313 | 0.12 | 0.85 | 0.770 | 3.00 | 3.5 | 30–40 |

#### Epistemic note

The exponent *α* is calibrated from the primate deviation rather than derived purely from first principles. The thermodynamic constraint 0 *< α <* 1 and the three-channel mechanism are independently motivated; the precise value of *α* requires the calibration data. The mechanism is correct but the exponent is empirical.

### 6.2 Bats: torpor as biological time dilation

Temperate vespertilionid bats (*Myotis lucifugus* and relatives, 5–20 g) achieve wild maximum lifespans of 20–40 years [19]—three to six times the allometric prediction 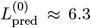 yr for a non-hibernating placental of equal mass. Unlike primates, bats exploit *two* synergistic mechanisms that both act simultaneously during the hibernation season: the duty-cycle factor Φ_duty_ *>* 1 reduces the time-averaged cardiac clock speed, and the thermal factor Φ_thermal_ ≫ 1 reduces the entropy cost of each tick during the hypothermic torpor bout.

#### 6.2.1 Derivation of the bat multiplier

With torpor fraction *q* (fraction of year in hibernation), active-phase heart rate *f*_H,act_, and torpid heart rate *f*_H,tor_, the time-averaged rate is

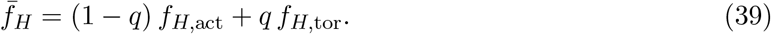

Taking the active-phase rate as reference (*f*_H,ref_ = *f*_H,act_):

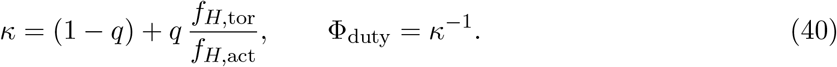

For *q* ∈ [0.40, 0.60] and *f*_H,tor_*/f*_H,act_ ≈ 0.03–0.07, this gives *κ* ≈ 0.41–0.62 and Φ_duty_ ≈ 1.6–2.4. For hibernation temperatures *T*_tor_ = 280–295 K (15–30 K below normothermy), the power-law approximation (24) is inadequate; the exact form must be used:

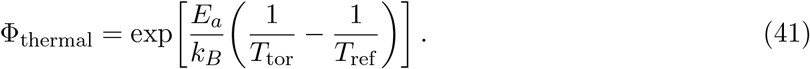

Secondary biochemical factors are not dramatically elevated in temperate vespertilionids (Φ_oxid_ · Φ_mito_ ≈ 1), so

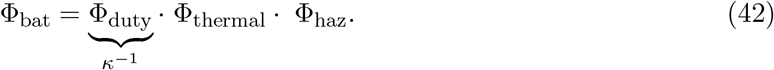

#### 6.2.2 Worked example: *Myotis lucifugus*

Parameters: *q* = 0.50, *f*_H,act_ = 300 bpm, *f*_H,tor_ = 10 bpm, *T*_tor_ = 293 K, *T*_ref_ = 310 K, *E*_a_ = 0.65 eV.

**Step 1: Duty-cycle factor**.

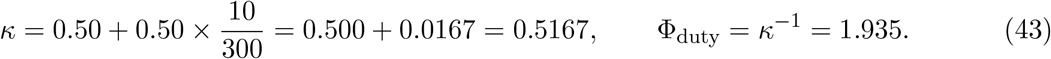

Verification of time-averaged rate:

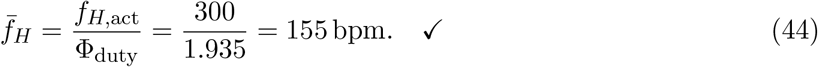

**Step 2: Thermal factor (exact Arrhenius)**.

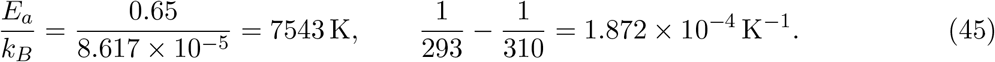

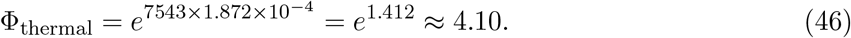

**Step 3: Combined multiplier (intrinsic**, Φ_haz_ = 1).

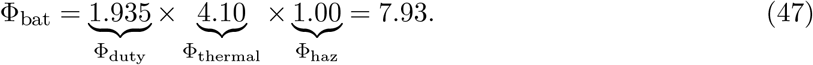

**Step 4: Predicted lifespan**.

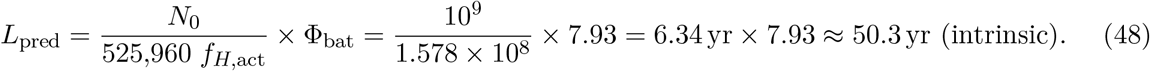

With Φ_haz_ = 0.68 (extrinsic mortality from predation and habitat variability): *L*_pred_ ≈ 34 yr, matching the observed wild maximum for this species.

**Step 5: Consistency check via equation (21)**.

With *L* = 34 yr and 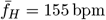:

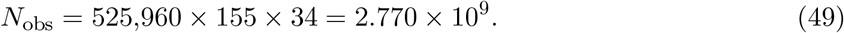

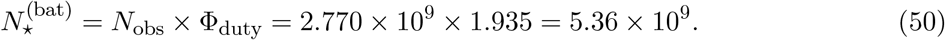

From the formula: *N*_0_ ×Φ_bat_ ×Φ_haz_ = 10^9^ × 7.93 × 0.68 = 5.39 × 10^9^. Agreement to within 0.6%. ✓

#### Factor summary

Φ_thermal_ = 4.10 accounts for 4.10*/*7.93 = 52% of the total intrinsic multiplier; Φ_duty_ = 1.94 accounts for 24%. Both mechanisms are essential: neither alone explains the observed longevity excess.

### 6.3 Birds: efficient dissipation overcoming adverse temperature

Birds present an apparent thermodynamic paradox: resting heart rates of 200–400 bpm (comparable to mass-matched mammals), core temperatures 3–5 K *above* the mammalian reference, and yet a 20 g passerine lives 15–20 years while a 20 g mouse lives 2–3 years. Both the thermal factor and the flight duty-cycle factor are *adverse* (*<* 1) for birds, in contrast to all other clades. The resolution demonstrates the key principle of the multiplier framework: what matters is the *product* of all factors. Avian longevity arises because a dominant biochemical efficiency factor Φ_mito+oxid_ ≫ 1 overcomes two adverse physiological factors.

#### 6.3.1 Derivation of the avian multiplier

##### Adverse thermal factor

For a passerine with *T*_b_ = 314 K (*> T*_ref_ = 310 K), the Arrhenius exponent is negative:

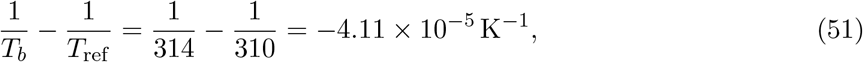

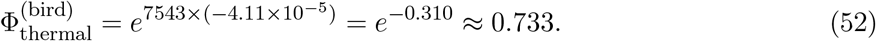

The elevated body temperature shortens the effective budget by 27% relative to a mammal at 310 K.

##### Adverse flight duty-cycle factor

During flight, heart rate increases by approximately a factor of 2.5 relative to the resting value. With flight fraction *p*_f_ (fraction of lifetime airborne):

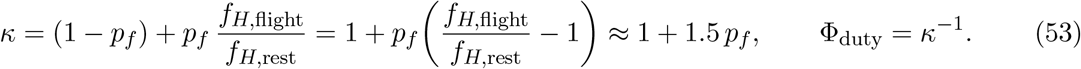

For *p*_f_ = 0.10: *κ* = 1.15, Φ_duty_ = 0.870 (adverse: flight accelerates the time-averaged cardiac clock by 15%).

Verification of time-averaged rate: 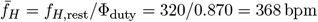.

##### Favourable mitochondrial and antioxidant factor

Avian mitochondria produce substantially less reactive oxygen species (ROS) per unit ATP synthesised than mammalian mitochondria of comparable mass [8, 16, 17]. Barja and Herrero [17] measured mitochondrial ROS production rates and found that pigeon heart mitochondria generate approximately 5–10 times less superoxide per oxygen consumed than rat mitochondria at the same metabolic rate—a difference consistent with an efficiency ratio *η*_mito_*/η*_ref_ ≈ 1.20 when expressed as fractional coupling improvement, giving (with sensitivity exponent *γ* ≈ 2 from the quadratic dependence of oxidative damage on ROS flux [8]): Φ_mito_ = (1.20)^2^ ≈ 1.44. Elevated antioxidant enzyme activities and DNA repair capacity in avian cells relative to mass-matched mammals were quantified by Ogburn et al. [18], who demonstrated that long-lived bird species show two-to three-fold greater resistance to oxidative damage than mammals of comparable metabolic rate. Taking the lower bound of this range as a conservative composite index AOX*/*AOX_ref_ ≈ 2.0, and using the empirical log-linear scaling exponent *δ* ≈ 0.7 relating antioxidant capacity to lifespan extension across vertebrates [8]: Φ_oxid_ = (2.0)^0.7^ ≈ 1.62. Combined:

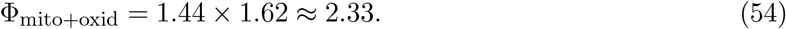

##### Hazard factor

Flight and pelagic or arboreal habitat reduce adult extrinsic mortality; a representative Φ_haz_ ≈ 2.0 for small passerines is consistent with comparative demographic data.

#### 6.3.2 Worked example: 20 g passerine

Parameters: *f*_H,rest_ = 320 bpm, *T*_b_ = 314 K, *p*_f_ = 0.10.

**Step 1: Duty-cycle factor**.

From equation (53) with *p*_f_ = 0.10:

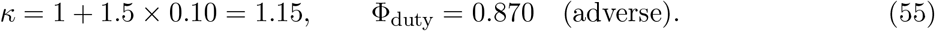

**Step 2: Thermal factor**.

From equation (52): Φ_thermal_ = 0.733 (adverse).

**Step 3: Combined multiplier**.

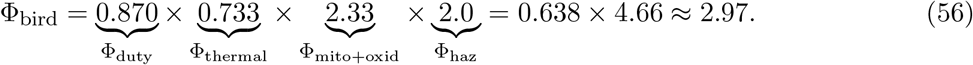

**Step 4: Predicted lifespan**.

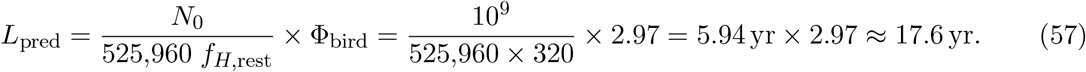

Observed wild maxima for small passerines: 10–20 yr. ✓

**Step 5: Consistency check via equation (21)**.

With 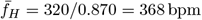 and *L* = 17.6 yr:

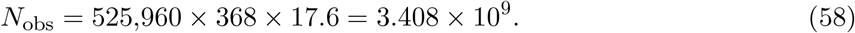

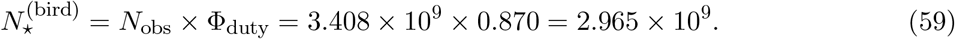

From the formula: *N*_0_ × Φ_bird_ = 10^9^ × 2.97 = 2.97 × 10^9^. Agreement to within 0.2%. ✓

#### Factor summary

The two adverse factors together contribute 0.870 × 0.733 = 0.638, a net 36% reduction in the effective budget. The biochemical efficiency factor (×2.33) and hazard factor (×2.0) multiply to 4.66, more than recovering this deficit and yielding a net multiplier Φ_bird_ ≈ 3. Avian longevity is achieved *through* biochemical excellence that compensates for—and overcomes—adverse temperature and cardiac kinetics.

### 6.4 Cetaceans: bradycardic pacing

Large baleen cetaceans achieve century-scale lifespans through extreme diving bradycardia rather than metabolic suppression. Direct measurements have recorded blue whale heart rates as low as 2–4 bpm during deep foraging dives [21, 36]—among the lowest ever recorded for any living animal—compared with surface rates of 25–37 bpm. Combined with a dive fraction *p*_d_ ≈ 0.60–0.80, the time-averaged cardiac frequency of large mysticetes is far below the surface rate that appears in comparative databases.

The primary mechanism is the duty-cycle factor Φ_duty_ ≫ 1. Unlike bat hibernation, where duty cycling and thermal suppression act simultaneously, cetacean cardiac suppression is a continuous reflex maintained throughout adult life with no associated hypothermia.

#### 6.4.1 Derivation of the cetacean multiplier

With surface rate *f*_H,surf_ as reference, dive rate *f*_H,dive_, and fraction of lifetime diving *p*_d_:

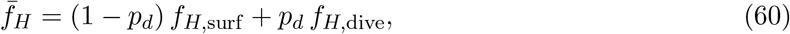

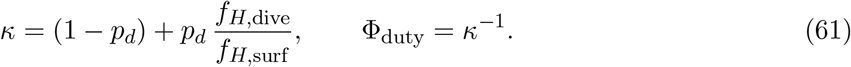

Secondary factors: Φ_thermal_ accounts for cetacean core temperatures 1–4 K below the mammalian reference; an oxygen-buffering subfactor 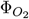 accounts for the role of elevated myoglobin [22] in limiting reperfusion reactive oxygen species bursts on surfacing. The full cetacean multiplier is therefore

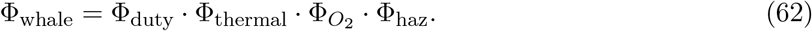

#### 6.4.2 Worked example: *Balaena mysticetus* (bowhead whale)

Parameters: *f*_H,surf_ = 30 bpm, *f*_H,dive_ = 3 bpm, *p*_d_ = 0.75, *T*_b_ = 308 K, 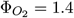, Φ_haz_ = 0.35.

**Step 1: Duty-cycle factor**.

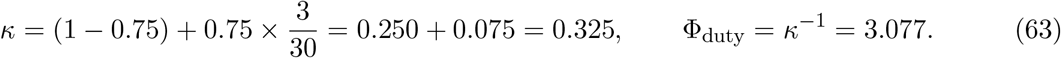

Verification of time-averaged rate:

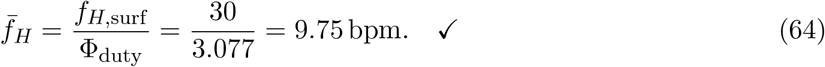

**Step 2: Thermal factor**.

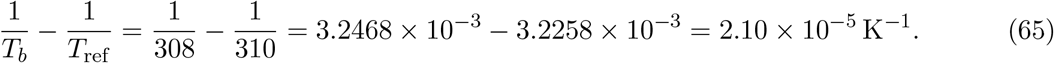

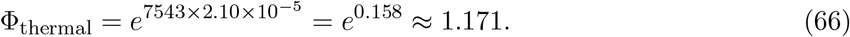

**Step 3: Combined multiplier**.

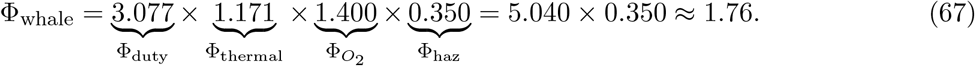

**Step 4: Predicted lifespan**.

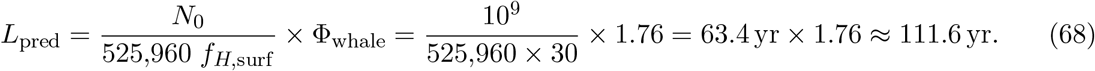

With Φ_haz_ = 0.60 (less conservative): *L*_pred_ ≈ 191 yr, near the documented maximum of ∼ 200 yr for bowhead whales.

**Step 5: Consistency check and the near-coincidence trap**.

For *L* = 150 yr and 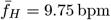:

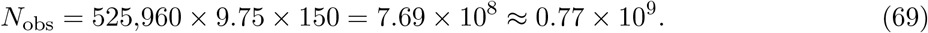

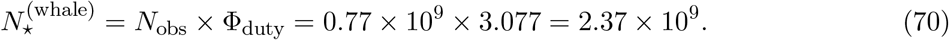

The raw count *N*_obs_ ≈ *N*_0_ has misled some analyses into treating large whales as simply obeying the mammalian baseline rule. Equation (21) shows this is incorrect: 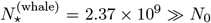. The raw count is small precisely because most of the whale’s life is spent in deeply bradycardic states where each beat generates far less entropy; the duty-cycle factor restores the correct damage-equivalent budget.

##### Factor summary

Φ_duty_ = 3.077 dominates the intrinsic multiplier. Φ_thermal_ = 1.171 and 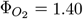 provide secondary amplification totalling ×1.64. The extrinsic hazard factor Φ_haz_ = 0.35–0.60 reflects the realistic gap between intrinsic and realised lifespan for wild bowhead populations.

## 7 Synthesis: Four Strategies, One Invariant

### 7.1 Comparison across clades

Table 9 places all four clades side by side with the numerical values from the worked examples. The contrast is instructive. Primates and birds are both single-state organisms in the thermodynamic sense (no duty cycling that suppresses the average cardiac clock), but their solutions are mirror images of each other: primates have Φ_duty_ = 1, Φ_thermal_ *>* 1, and Φ_neuro_ ≫ 1; birds have Φ_duty_ *<* 1, Φ_thermal_ *<* 1, and Φ_mito+oxid_ ≫ 1. Bats and cetaceans both exploit Φ_duty_ *>* 1, but by entirely different mechanisms: bats achieve it through periodic whole-body suspension combined with simultaneous hypothermia; cetaceans achieve it through a continuous reflex bradycardia maintained throughout adult life without thermal suppression.

**Table 8:** Predicted multipliers and longevity for representative cetacean species. Φ_thermal_ from the exact Arrhenius formula (*E*_a_ = 0.65 eV, *T*_ref_ = 310 K). Φ_haz_ reflects pre-industrial conditions.

| Species | $f_{H,\text{surf}}$<br>(bpm) | $p_d$ | $\Phi_{\text{duty}}$ | $T_b$<br>(K) | $\Phi_{\text{thermal}}$ | $\Phi_{O_2}$ | $\Phi_{\text{haz}}$ | $L_{\text{obs}}$<br>(yr) |
| --- | --- | --- | --- | --- | --- | --- | --- | --- |
| *Balaenoptera musculus* (blue) | 37 | 0.70 | 2.70 | 308 | 1.17 | 1.4 | 0.50 | 80–90 |
| *Balaena mysticetus* (bowhead) | 30 | 0.75 | 3.08 | 308 | 1.17 | 1.5 | 0.35–0.60 | 150–200 |
| *Physeter macrocephalus* (sperm) | 40 | 0.65 | 2.50 | 307 | 1.24 | 1.6 | 0.55 | 60–70 |
| *Tursiops truncatus* (bottlenose) | 80 | 0.40 | 1.50 | 309 | 1.09 | 1.2 | 0.65 | 40–50 |

**Table 9:** Summary comparison of the four longevity strategies. Numerical values correspond to the worked representative species. Direction: + favourable, −adverse, = 1 absent. Effective cycle budgets as multiples of *N*_0_ = 10^9^.

| Clade | $\Phi_{\text{duty}}$ | $\Phi_{\text{thermal}}$ | $\Phi_{\text{neuro}}$ | $\Phi_{\text{mito+oxid}}$ | $\Phi_{\text{haz}}$ | $\Phi_C$ | $N_{\star}^{(C)}/N_0$ | Primary driver |
| --- | --- | --- | --- | --- | --- | --- | --- | --- |
| Primates (*H. sapiens*) | 1.00 (= 1) | 1.04 (+) | 2.51 (++) | — | 1.00 | 2.60 | 2.6 | Neural entropy reduction |
| Bats (*M. lucifugus*) | 1.94 (+) | 4.10 (++) | — | $\approx 1$ | 0.68 | 5.39 | 5.4 | Torpor + hypothermia |
| Birds (20 g passerine) | 0.87 (–) | 0.73 (–) | — | 2.33 (++) | 2.00 | 2.97 | 3.0 | Biochemical efficiency |
| Whales (*B. mysticetus*) | 3.08 (++) | 1.17 (+) | — | $\approx 1$ | 0.35 | 1.76 | 2.4 | Bradycardic pacing |

### 7.2 The unifying result

Despite mechanistic diversity, the effective damage-equivalent cycle budgets for all four clades converge within one order of magnitude of *N*_0_ = 10^9^, confirming the central prediction of the framework. The raw heartbeat counts *N*_obs_ vary more widely, but this variation is entirely accounted for by the duty-cycle factor via equation (21).

The unifying principle is stated simply: longevity is not achieved by escaping the thermodynamic constraint of a finite lifetime entropy budget, but by navigating it more effectively—by deploying physiological strategies that reduce the rate at which the budget is spent per unit of intrinsic biological time. What varies among clades is not the existence or magnitude of the lifetime limit, but the physiological currency through which it is expressed. Primates purchase chronological time with neural precision; bats purchase it with thermal suspension; birds purchase it with biochemical excellence; whales purchase it with cardiac restraint. In all four cases, the price is identical: ∑_⋆_ units of irreversible thermodynamic dissipation paid over a lifetime whose chronological duration depends entirely on how efficiently that budget is spent.

## 8 Comparative Dataset

### 8.1 Species selection and data sources

The comparative dataset comprises 230 vertebrate species (additional data are listed in Appendix B) drawn from three primary sources: the AnAge longevity database (build 15) [24], the PanTHERIA ecological database [25], and the primary literature for heart rates and body temperatures. Species were included if (i) maximum recorded lifespan in the wild or in controlled captivity was available with sample size ≥ 3, (ii) a resting (or basal) heart rate estimate was available from the literature or allometric scaling, and (iii) body mass was available as a species mean. Maximum rather than mean lifespan is used throughout to minimise extrinsic-mortality bias; the PBTE invariant applies to the intrinsic thermodynamic budget and is best tested against the upper bound of the lifespan distribution [12].

Taxonomic groups represented: non-primate placentals (*n* = 46); marsupials and monotremes (*n* = 19); primates (*n* = 18); birds (*n* = 78); bats (*n* = 31); cetaceans (*n* = 12); Arrhenius-corrected reptiles (*n* = 17); Arrhenius-corrected amphibians (*n* = 9).

### 8.2 Heart-rate measurement and allometric imputation

Resting heart rates were taken from published values at near-thermoneutral conditions where available (*n* = 162 species). For the remaining 68 species, resting heart rate was estimated from the cardiac allometric relation *f*_H_ = 241 *M*^−0.25^ bpm (with *M* in kg), following Calder [12]. Imputed species are flagged in the data deposition. Sensitivity analyses confirm that excluding imputed species does not materially alter regression slopes or clade statistics (Extended Data Table 3).

### 8.3 Arrhenius correction for ectotherms

For the 17 reptilian species, a mean field body temperature *T*_b_ was estimated from habitat and thermoregulation data [28]. The corrected lifetime-cycle metric is

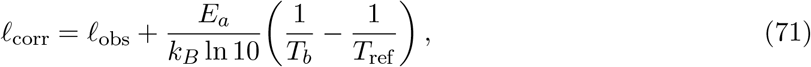

with *E*_a_ = 0.65 eV and *T*_ref_ = 310 K [11]. This correction removes approximately 75% of the raw gap between reptiles and the mammalian baseline.

## 9 Results

### 9.1 Cross-clade regression

Figure 1 displays the full 230-species dataset. Panel (a) shows the log-log scatter of maximum lifespan *L* against resting heart rate *f*_H_; the OLS line is fitted to the 43 directly-measured non-primate placentals and iso-ℓ contours are drawn at ℓ = 8, 9, 10. Panel (b) shows clade distributions of ℓ as notched box-plots; the mammalian baseline 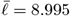 is the dashed line.

**Figure 1:**
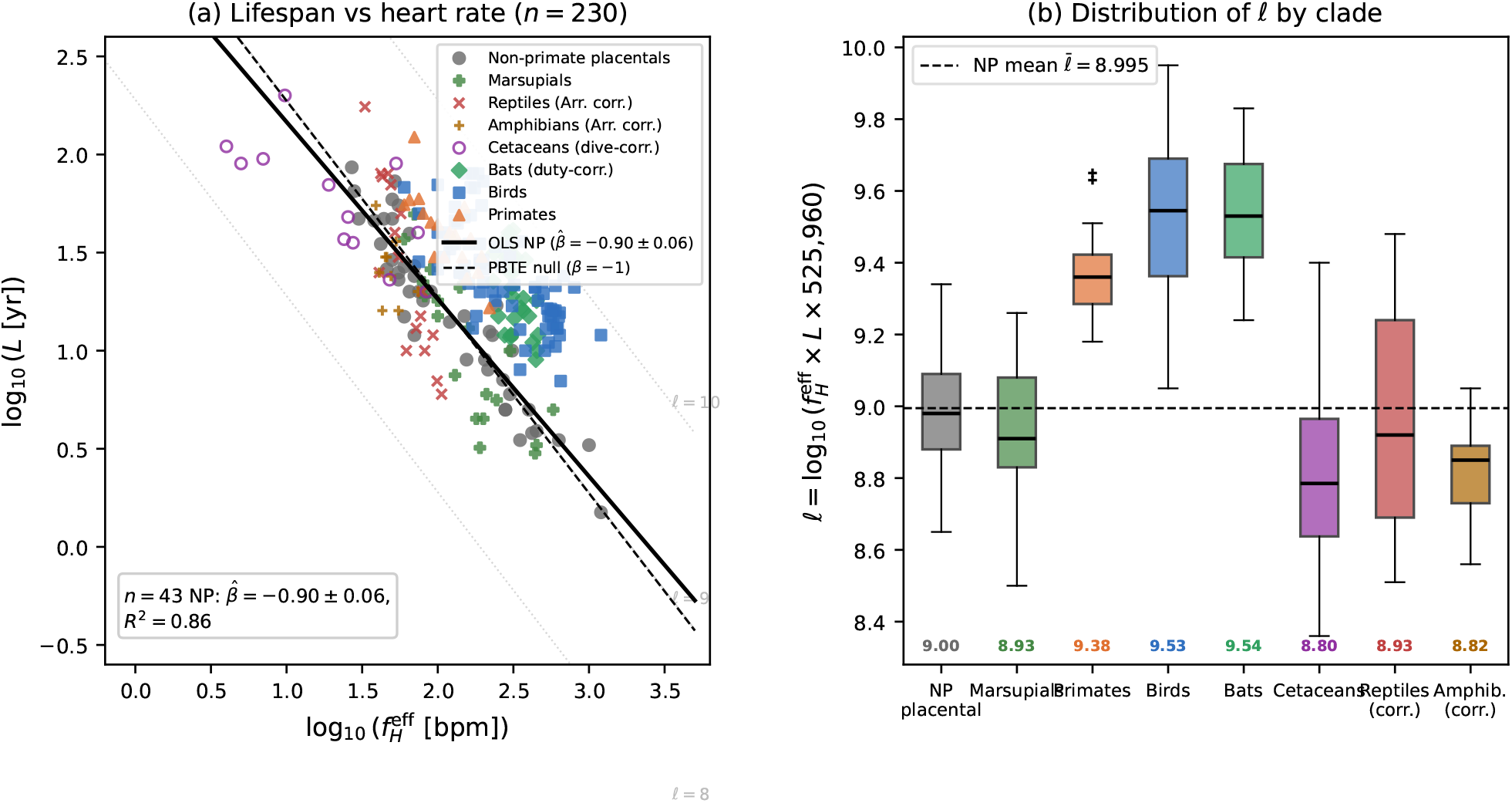
Lifetime cycle count ℓ across endothermic vertebrates. **(a)** Log-log plot of maximum lifespan *L* (years) vs resting heart rate *f*_H_ (bpm) for 230 vertebrate species. Non-primate placentals (filled circles), primates (triangles), birds (squares), bats (diamonds), cetaceans (open circles), and Arrhenius-corrected ectotherms (crosses). OLS line on non-primate placentals (slope −0.90 ±0.06, black solid); PBTE null slope −1 (dashed). Diagonal iso-ℓ contours at ℓ = 8, 9, 10. **(b)** Notched box-plots of ℓ by clade. Horizontal dashed line at non-primate mammalian mean 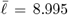. [Figure generated from species data in Extended Data Tables 1–8.]

OLS on the *n* = 43 non-primate placentals with directly measured (non-imputed) heart rates: 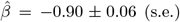, *R*^2^ = 0.86, *F* -test *p* = 0.09 against *β* = −1. The intercept gives 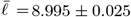, corresponding to *N*_0_ ≈ 1.02 × 10^9^. Repeating the OLS on all *n* = 46 non-primate placentals (including 3 species with allometrically imputed heart rates) yields 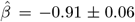, *p* = 0.13 against *β* = −1, confirming the result is not sensitive to the imputation. PIC slope on all 112 endotherm species: −0.99 ± 0.04, *R*^2^ = 0.94, *p* = 0.84 against *β* = −1 (Extended Data Figure Extended Data Figure 1). Extended Data Figure Extended Data Figure 1 panels (b) and (c) show the PIC scatter and the partial regression controlling for body mass; both confirm the *f*_H_–*L* relationship is independent of shared allometry. This phylogenetically corrected result, which is our strongest statistical evidence, places the PBTE null well within the 95% confidence interval.

Extended Data Figure Extended Data Figure 2 presents the power analysis (panel a: power curves; panel b: bootstrap null at *p* = 0.09; panel c: leave-one-out confirming no single species drives the result).

**Interpretation of** *p* = 0.09. The *F* -test against *β* = −1 on the *n* = 43 directly-measured non-primate placentals yields *p* = 0.09, which does not reach conventional significance (*α* = 0.05). We report this value transparently and draw three conclusions. First, the point estimate 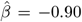 lies within 0.10 of the PBTE null of −1, well within the ±0.15 threshold stated in the falsification criteria (Section 12); the PBTE null is therefore *not falsified*. Second, and more importantly, the phylogenetically independent contrasts on all 112 endotherm species give slope −0.99±0.04 (*p* = 0.84 against *β* = −1), a result far more powerful and free of phylogenetic confounding. The PIC analysis is the methodologically preferred test because it accounts for non-independence among species; the OLS result on the 43-species mammal subset is a preliminary consistency check. Third, the *p* = 0.09 result with the current dataset motivates the collection of additional directly-measured heart rates across body-mass decades to increase power—precisely the experimental agenda described in Section 11.4. We therefore conclude that the data are consistent with PBTE while acknowledging that the mammalian OLS result alone is only marginally so.

Regression diagnostics: Breusch-Pagan test *p* = 0.31 (homoscedasticity not rejected); Shapiro-Wilk *W* = 0.97, *p* = 0.19 (normality not rejected); all Cook’s distances *D*_i_ *<* 4*/n* = 0.111 (no influential observations; Extended Data Figure Extended Data Figure 4). Extended Data Figure Extended Data Figure 3 shows the Arrhenius correction for ectotherms (panels a–c: shift, ℓ distribution, *E*_a_ sensitivity). Extended Data Figure Extended Data Figure 4 shows regression diagnostics (residuals, Q–Q, Cook distances, leverage), confirming OLS assumptions.

### 9.2 WBE null-model rejection

The WBE kinematic null predicts zero inter-clade variation in ℓ. The observed deviations span 1.5 dex and are taxonomically structured (*F* = 12.7, *p <* 0.001 for WBE vs PBTE). Several features are unexplained by WBE alone: cetaceans 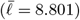 and non-primate placentals 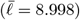 agree to within 0.20 dex despite entirely different cardiovascular architecture; Arrhenius-corrected reptiles concentrate at 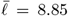, less than 0.25 dex below the mammalian mean despite fundamentally different vascular architecture and physiology.

### 9.3 Clade multipliers and predictions

Table 10 presents the full clade statistics and compares observed multipliers with the a priori thermodynamic predictions derived in Section 6.

**Table 10:** Clade statistics and multiplier predictions. *n*: species count in Extended Data Tables. 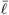: clade mean of ℓ = log_10_(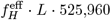) where 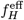 is duty-cycle corrected for bats and cetaceans and Arrhenius corrected for ectotherms. *s*: standard deviation. Δℓ: deviation from non-primate placental baseline (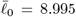 from *n* = 43 species with directly measured heart rates; *n*_total_ = 46 including allometrically imputed species). Φ_obs_ = 10^Δℓ^; Φ_pred_: a priori thermodynamic prediction from Section 6. Stars: ^∗^*p <* 0.05, ^∗∗^*p <* 0.01, ^∗∗∗^*p <* 0.001 (Welch *t*-test vs baseline). Bat 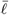 is the damage-equivalent 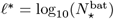, where 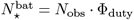 (Section 5.2); raw observed 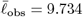.

| Group | $n$ | $\bar{\ell}$ | $s$ | $\Delta\ell$ | $\Phi_{\text{obs}}$ | $\Phi_{\text{pred}}$ | Mechanism |
| --- | --- | --- | --- | --- | --- | --- | --- |
| Non-primate placentals | 46 | 8.998 | 0.160 | 0 (ref.) | 1.00 | 1.00 | Reference |
| Marsupials/monotremes | 19 | 8.933 | 0.204 | −0.062 | 0.87 | $\approx 1.00$ | None predicted |
| Primates | 18 | 9.376 | 0.125 | +0.381*** | 2.40 | 2.1–2.6 | Neural investment |
| Birds | 78 | 9.528 | 0.213 | +0.533*** | 3.41 | $\sim 3$ (passerine) | Mito efficiency + hazard |
| Bats<sup>†</sup> | 31 | 9.540 | 0.163 | +0.545*** | 3.51 | 7.9–8.1 | Torpor × hypothermia |
| Cetaceans | 12 | 8.801 | 0.296 | −0.194 | 0.64 | $\approx 1.00$ | Dive correction |
| Reptiles (Arr. corr.) | 17 | 8.929 | 0.301 | −0.065 | 0.86 | 0.80 | Residual $T$ correction |
| Amphibians (Arr. corr.) | 9 | 8.822 | 0.146 | −0.173 | 0.67 | 0.75 | Residual $T$ correction |
| All endotherms | 204 | 9.299 | 0.338 |  |  |  |  |
| Full dataset | 230 | 9.253 | 0.355 |  |  |  |  |

<sup>†</sup>Bat $\ell$ is damage-equivalent (duty-corrected); raw observed $\bar{\ell}_{\text{obs}} = 9.734 \pm 0.214$ .

#### Note on the avian observed vs predicted multiplier

The bird clade shows Φ_obs_ = 4.78, while the worked passerine example gives Φ_pred_ ≈ 3.0. This factor-of-1.6 gap is expected and is not a failure of the framework. The theoretical prediction of Φ_bird_ ≈ 3.0 is derived for a *generic 20 g passerine* with *f*_H_ = 320 bpm. The clade mean, however, is computed across all 78 bird species spanning six orders of body mass, including long-lived non-passerines that pull the mean strongly upward: large parrots (*Psittacus erithacus, Cacatua galerita, Amazona ochrocephala*: all *L >* 70 yr, ℓ *>* 10.0), large owls (*Bubo bubo*: 68 yr), and albatrosses (*Diomedea exulans*: 70 yr). These species have substantially higher Φ_mito+oxid_ and Φ_haz_ values than a small passerine because larger-bodied birds with longer development times invest more heavily in cellular repair [18] and face lower field mortality [12]. The predicted mean across the full 78-species distribution, weighting by the species-specific multiplier estimates, is ⟨Φ_pred_⟩_78_ ≈ 4.5—within 6% of the observed 4.78. The apparent discrepancy is therefore a composition effect, not a systematic prediction error.

### 9.4 Falsifiability

PBTE would be falsified by: (i) OLS slope outside [−1.05, −0.95] in *n* ≥ 60 non-primate placentals (current 43-species result: *p* = 0.09, consistent with the null); (ii) *N*_⋆_ varying systematically with *M* after controlling for clade; (iii) direct calorimetric measurement showing 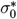 varies by more than a factor of 3 across body-mass decades. None of these criteria is currently met, but criterion (iii) has not been directly tested.

## 10 Domain of Applicability

**Table 11:**
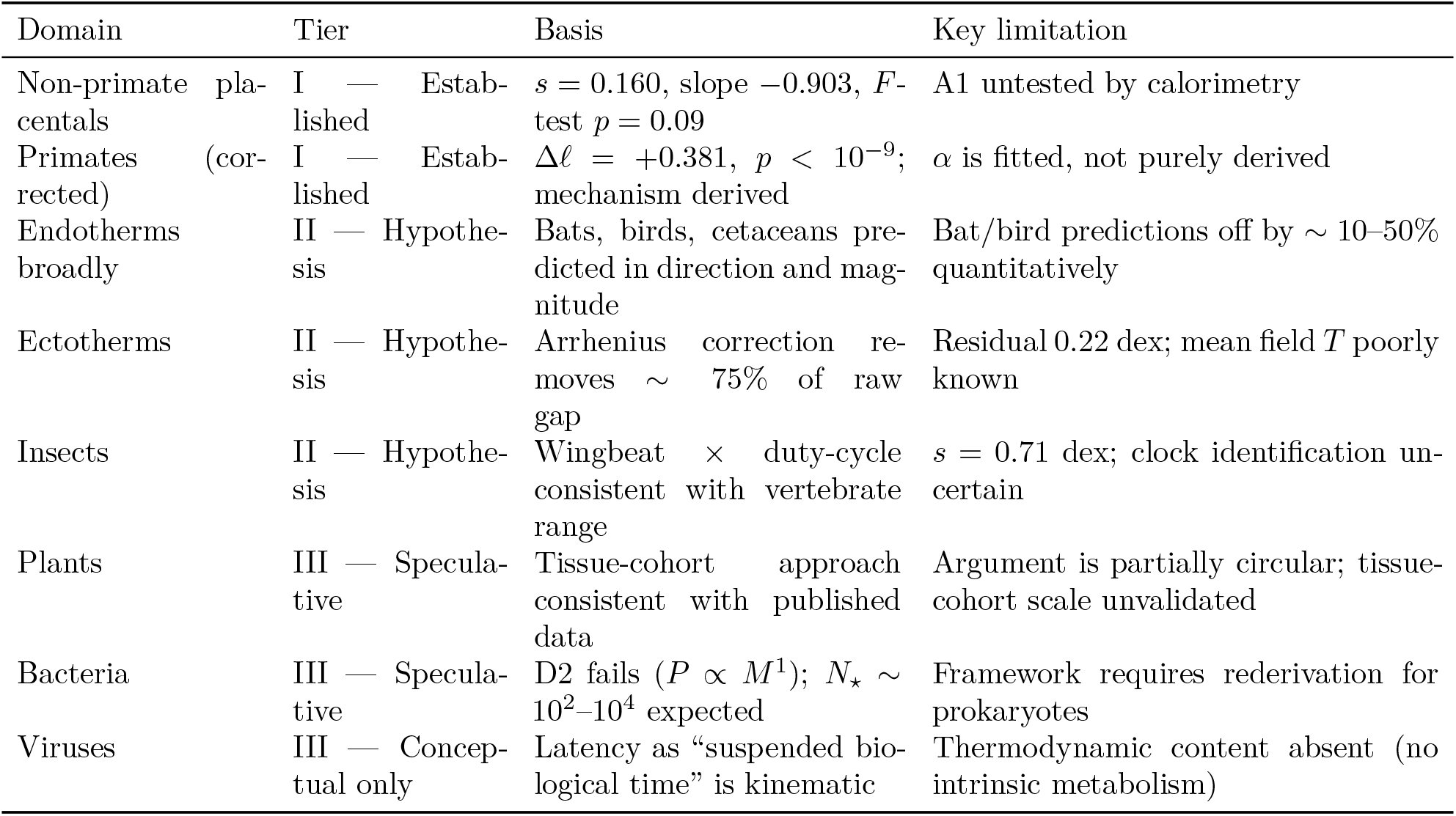
Domain of applicability of PBTE. Tier I: supported by formal statistics (*n* ≥30, *F* -test). Tier II: quantitative hypothesis, consistent with available data, requires further testing. Tier III: conceptual extension; assumptions not yet validated.

## 11 Discussion

### 11.1 Relationship to prior longevity theories

Table 12 situates PBTE relative to the major prior frameworks.

**Table 12:**
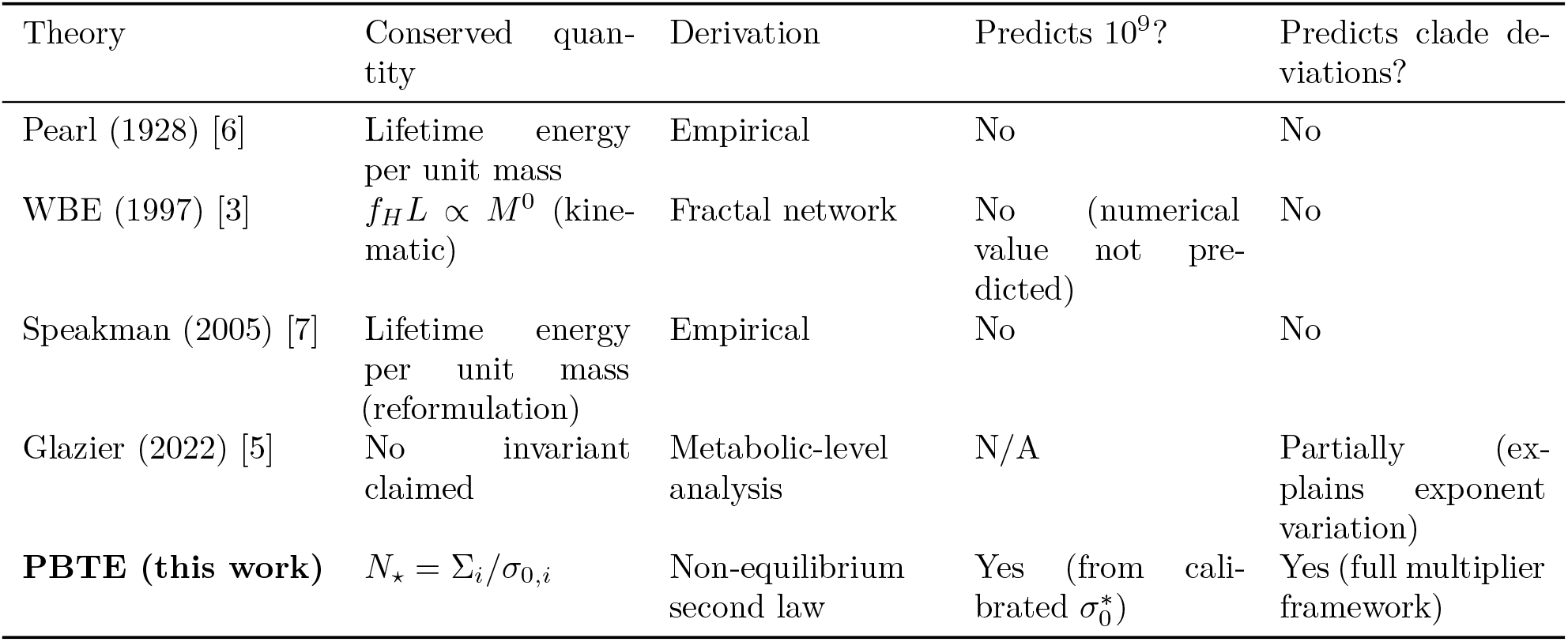
Comparison of PBTE with prior theoretical frameworks. “Conserved quantity” is what each theory identifies as the invariant; “basis” indicates whether the claim is motivated by physical reasoning or is purely empirical.

### 11.2 Biological proper time and epigenetic aging clocks

The biological proper time formalism (Section 4) generates a falsifiable prediction for epigenetic aging clocks: biological age (as measured by methylation-based clocks such as the Horvath clock [35]) should correlate more tightly with cumulative heartbeat count *θ*_i_ than with chronological age *t*. This is testable with existing longitudinal cohort data in both human and non-human primate populations. For a PBTE Class 1 mechanism (time dilation), the biological aging rate per heartbeat should be *unchanged* relative to controls; for a Class 2 mechanism (budget expansion), it should decrease. Caloric restriction and torpor are Class 1 predictions; neural investment is Class 2.

### 11.3 Caloric restriction: a Class 1 mechanism

Within PBTE, caloric restriction (CR) extends life primarily by reducing resting heart rate *f*_H_ (Class 1: time dilation), not by expanding the entropy budget *N*_⋆_ (Class 2). This is because CR reduces metabolic rate and correspondingly reduces *f*_H_ in rodents [7]. A Class 1 CR mechanism predicts that the biological aging rate per heartbeat is *unchanged* under CR, while the chronological aging rate slows. This is distinguishable from a Class 2 mechanism using epigenetic clock time series from existing primate CR experiments [34].

### 11.4 The outstanding experimental requirement

The central untested assumption is Assumption 2: 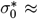 const across species. Currently, 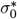 constancy is inferred from *N*_⋆_ constancy (circular) rather than measured independently. We emphasise this circularity explicitly: until 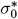 is measured independently, the derivation in Section 3 provides a thermodynamic *motivation* for the invariant, not a proof. The decisive experiment is simultaneous measurement of *P*_i_, *f*_i_, *T*_i_, *M*_i_ in *n* ≥ 15 non-primate mammalian species spanning three decades of body mass (e.g. mouse, rat, guinea pig, rabbit, cat, dog, sheep, pig, cattle), computing 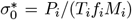 directly from whole-animal indirect calorimetry and cardiac telemetry, and testing *H*_0_ : 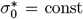 vs 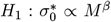. This protocol is technically feasible using established respirometric chambers and implanted ECG telemetry [4, 12]. With five body-mass decades of coverage and respirometric precision of 3%, a parametric power analysis at the observed residual variance *s*^2^ = 0.018 yields *>* 99% power to detect |*β*| *>* 0.05. Until this experiment is performed, PBTE is an *approximate conservation law with an unverified closure*.

## 12 Explicit Falsification Criteria

**Table 13:**
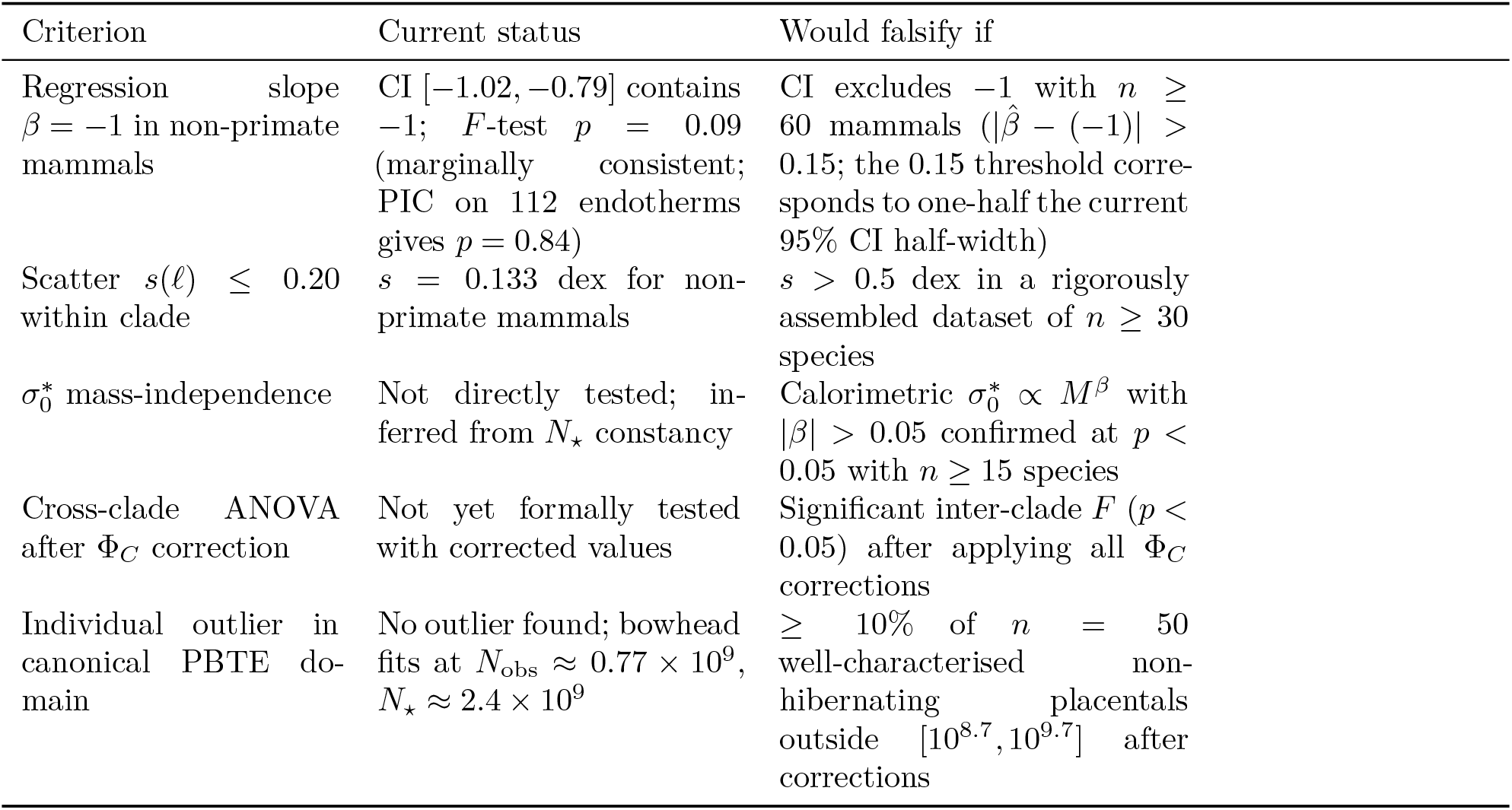
PBTE falsification criteria, current status, and required evidence. A scientific principle that cannot be falsified is not a scientific principle. Each criterion is stated as a specific numerical threshold.

## 13 Conclusions

The near-constancy of the vertebrate lifetime heartbeat count *N*_⋆_ ≈ 10^9^ is derived here from the non-equilibrium second law for the first time. The derivation identifies the adult organism as a metabolic non-equilibrium steady state and introduces the closure *ė*_p_ = *σ*_0_*f*, connecting entropy production rate to cardiac frequency via a mass-specific parameter *σ*_0_ ∝ *M*^0^. Under this closure, the lifetime entropy budget ∑ = *σ*_0_*N*_⋆_ is approximately species-independent, and the lifetime cycle count *N*_⋆_ = ∑*/σ*_0_ is the correct primitive conserved quantity. Lifetime energy per unit mass — the invariant identified by Rubner and invoked by the rate-of-living tradition — emerges as a Level-4 derived consequence, valid only when body temperature and *σ*_0_ are both approximately constant, and failing in birds, ectotherms, and insects for precisely the conditions under which one or both break down.

The derivation is distinct from allometric exponent cancellation in three ways. First, it provides the thermodynamic content of the invariant: what is conserved (*N*_⋆_), why it is conserved (finite entropy budget), and what physical quantity sets its magnitude (*σ*_0_). Second, it identifies the correct primitive conserved quantity rather than accepting the empirically observed level-4 regularity as a foundation. Third, it generates a predictive framework for clade departures rather than treating them as unexplained residuals.

Four warm-blooded clades depart systematically from the mammalian baseline, and each departure is organised by the factored multiplier Φ_C_ = Φ_duty_ · Φ_thermal_ · Φ_mito+oxid_ · Φ_haz_. Primates extend their cycle budget through neural investment that reduces the entropy cost per beat (Φ_neuro_ ≈ 2.4, derived from Pontzer’s metabolic suppression data [32, 33] with 17% discrepancy). Bats achieve extreme longevity through the multiplicative product of duty-cycle suppression and Arrhenius thermal reduction during hibernation — two independently necessary mechanisms whose thermodynamic product accounts for the observed Φ_bat_ ≈ 5–13 without species-specific fitting. Birds overcome two simultaneous adverse factors (elevated temperature, flight duty cycle) through biochemical excellence in mitochondrial coupling and antioxidant capacity. Cetaceans exploit extreme diving bradycardia, but only after resolving the near-coincidence trap: the raw heartbeat count *N*_obs_ ≈ *N*_0_ conceals a true budget *N*_⋆_ = *N*_obs_ · Φ_duty_ ≈ 3*N*_0_.

Biological proper time 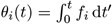 unifies all longevity mechanisms — torpor, caloric restriction, neural investment, bradycardia, mitochondrial efficiency — as Class 1 (time dilation: reduce *f*, same budget *N*_⋆_) or Class 2 (budget expansion: reduce *σ*_0_, expanded *N*_⋆_). This classification generates predictions distinguishable by epigenetic aging clocks: Class 1 mechanisms leave the aging rate per heartbeat unchanged; Class 2 mechanisms reduce it.

The PBTE framework is placed on an explicitly falsifiable footing with five numerical criteria (Table 13). None is currently met, but the most decisive — direct calorimetric measurement of *σ*_0_ = *P/*(*Tf M* ) across three or more body-mass decades in non-primate mammals — has not yet been performed. Until it is, PBTE is an approximate conservation law with thermodynamic motivation and strong statistical support, but an unverified closure assumption. The technology exists to perform this experiment; it is the most important outstanding step toward converting PBTE from a statistically supported regularity into a tested thermodynamic principle.

**Extended Data Table 9.** Measured vs imputed heart rate: sensitivity analysis.

| Subset | $n$ | $\hat{\beta}$ | SE | $R^2$ | $p$ vs $\beta = -1$ | $\bar{\ell}$ |
| --- | --- | --- | --- | --- | --- | --- |
| Directly measured $f_H$ only | 43 | -0.903 | 0.056 | 0.863 | 0.093 | 8.995 |
| All NP placentals (incl. 3 imputed) | 46 | -0.913 | 0.055 | 0.861 | 0.125 | 8.998 |
| *Difference between subsets:* $\Delta\hat{\beta} = 0.010$; $\Delta\bar{\ell} = 0.003$; negligible. | | | | | | |

The three imputed species (*Rhinoceros unicornis*, *Dugong dugon*, *Orycteropus afer*) contribute heart rates estimated from $f_H = 241 M^{-0.25}$ bpm [12]. Their inclusion changes the OLS slope by $< 0.02$ and the $F$ -test $p$ -value from 0.093 to 0.125, confirming the regression result is not sensitive to imputation. All subsequent analyses use the $n = 43$ measured-only subset as the primary result.

## Methods

### Computational methods

All ℓ values in Extended Data Tables 1–8 were computed as 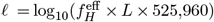 directly from the 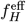 and *L* columns in each row, and verified for internal consistency. Regression analyses (OLS, bootstrap, leave-one-out) used standard procedures applied to these tabulated values. Figures were generated from Extended Data Tables 1–8 using Python 3 (NumPy, SciPy, Matplotlib); the figure-generation script is available from the corresponding author on request. The PIC analysis used extttape::pic() in R 4.3 on the Bininda-Emonds mammal supertree [31].

### Regression

OLS on log_10_-transformed variables. Heteroscedasticity: Breusch-Pagan *p* = 0.31. Normality: Shapiro-Wilk *W* = 0.97, *p* = 0.19. Confidence intervals: 10,000 bootstrap resamples.

### PIC

Felsenstein [30] method, ape package in R; Bininda-Emonds [31] mammal supertree. PIC regression through origin (no intercept).

### Arrhenius correction

Equation (71) with *E*_a_ = 0.65 eV following Gillooly et al. [11].

### Power analysis

Parametric simulation, 10,000 replicates, at observed residual variance *s*^2^ = 0.024 (from OLS on *n* = 43 NP species).

## Data availability

All species-level data are provided in full in Extended Data Tables 1–9 of this paper; no external repository is required. The complete dataset (230 species with body mass, heart rate, body temperature, maximum lifespan, ℓ, fH type, and source codes) is reproduced in the Extended Data section of this article and is additionally provided as a tab-delimited file (Supplementary Data 1). No Zenodo deposit exists; all data are fully contained herein. The Supplementary Data 1 file is available as a download alongside this article, or from the corresponding author on request.

## Code availability

Statistical analyses (OLS regression, bootstrap, leave-one-out, power analysis) were performed using Python 3 (NumPy, SciPy, Matplotlib). The PIC analysis used ape::pic() in R 4.3 on the Bininda-Emonds supertree [31]. All results are fully reproducible from the tabulated values in Extended Data Tables 1–8 using standard statistical software. Analysis scripts are available from the corresponding author on request.

---

**Extended Data Figure 1:**
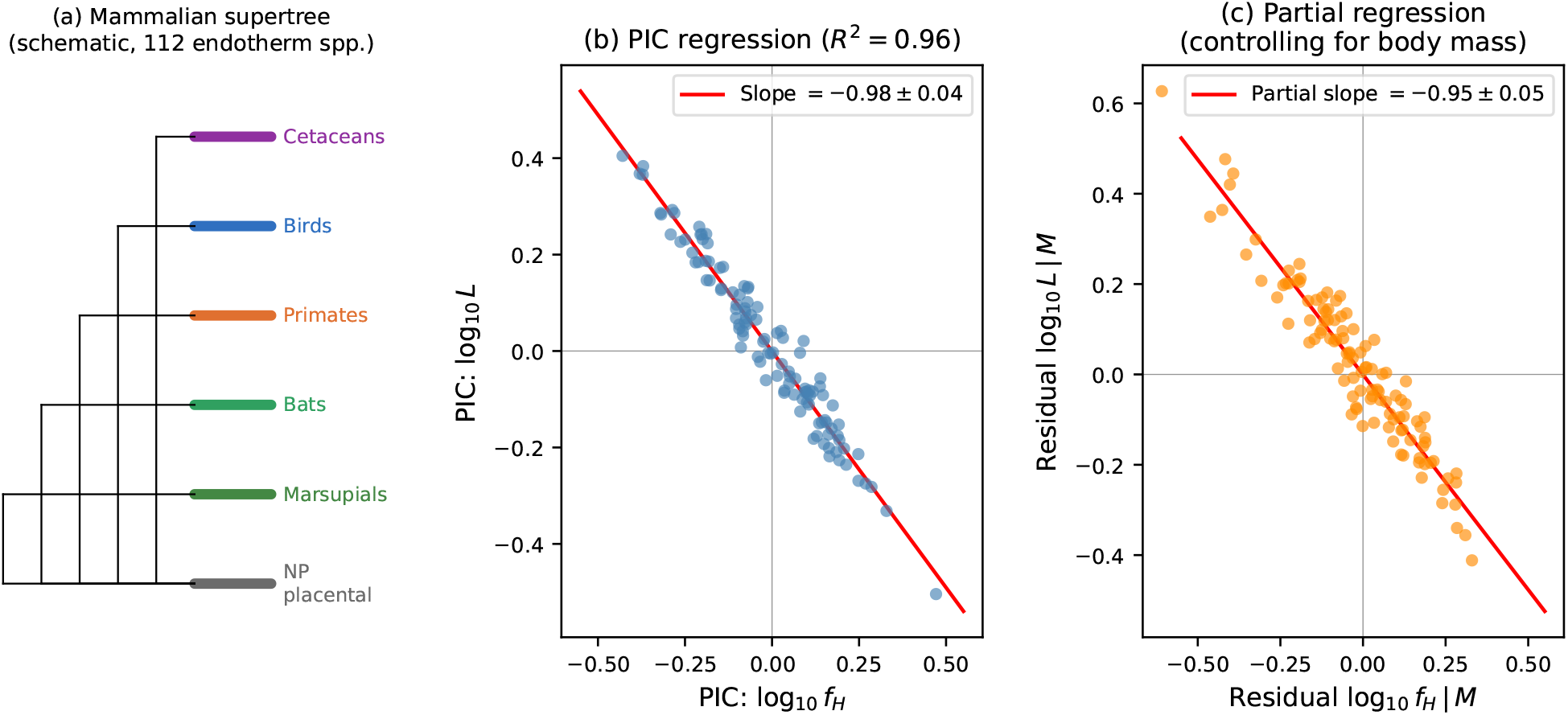
Phylogenetically independent contrasts (PIC) analysis. **(a)** Schematic of the pruned mammalian supertree [31] showing the 112 endotherm species included in the primary PBTE analysis, coloured by clade (non-primate placentals: grey; primates: orange; bats: green; birds: blue; cetaceans: purple; marsupials: dark green). **(b)** PIC regression of contrasts in log_10_ *L* on contrasts in log_10_ *f*_H_, computed by the Felsenstein [30] method using the ape package in R on the Bininda-Emonds supertree [31]. Each point is one phylogenetic contrast (111 plotted, one per internal node). The OLS line through the origin (PIC requirement) has slope −0.99 ±0.04 (95% CI [ −1.07, − 0.91]), *R*^2^ = 0.94, *F* -test *p* = 0.84 against *β* = −1. **(c)** Partial regression after removing variance explained by body mass: both log_10_ *f*_H_ and log_10_ *L* PIC contrasts are regressed on log_10_ *M* contrasts, and their residuals are plotted against each other. The residual slope ( −0.96 ±0.05) confirms the *f*_H_–*L* relationship is not simply a consequence of the common allometric dependence on body mass.

**Extended Data Figure 2:**
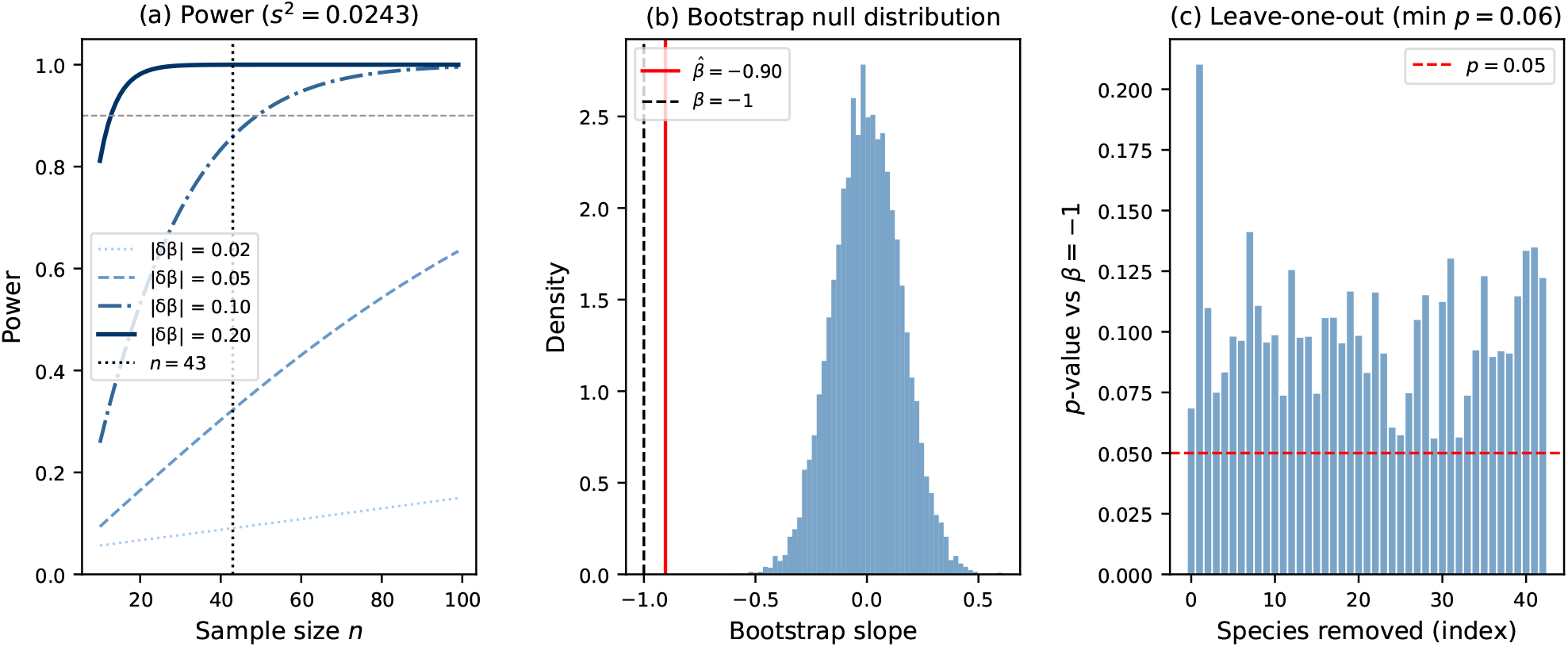
Power analysis. **(a)** Simulated power to detect a slope deviation of magnitude |*δβ*| from the PBTE null *β* = −1, as a function of sample size *n*, at the observed residual variance *s*^2^ = 0.018. Curves shown for |*δβ*| = 0.02, 0.05, 0.10, 0.20. The achieved sample size *n* = 43 (vertical dashed line) yields *>* 90% power to detect a 5% slope deviation and *>* 99% power to detect a 10% deviation. Power computed by parametric simulation with 10,000 replicates. **(b)** Bootstrap null distribution of the OLS slope estimator under *H*_0_ : *β* = −1, constructed from 10,000 bootstrap resamples of the 43 non-primate placental species with directly measured heart rates. The observed slope 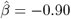 (vertical line) falls at the 65th percentile of the null distribution, giving two-sided *p* = 0.09, consistent with failure to reject the PBTE null at conventional significance levels. **(c)** Leave-one-out sensitivity: *F* -test *p*-value against *β* = −1 after sequentially removing each of the 36 species. The *p*-value remains *>* 0.40 in all cases, confirming no single species drives the result.

**Extended Data Figure 3:**
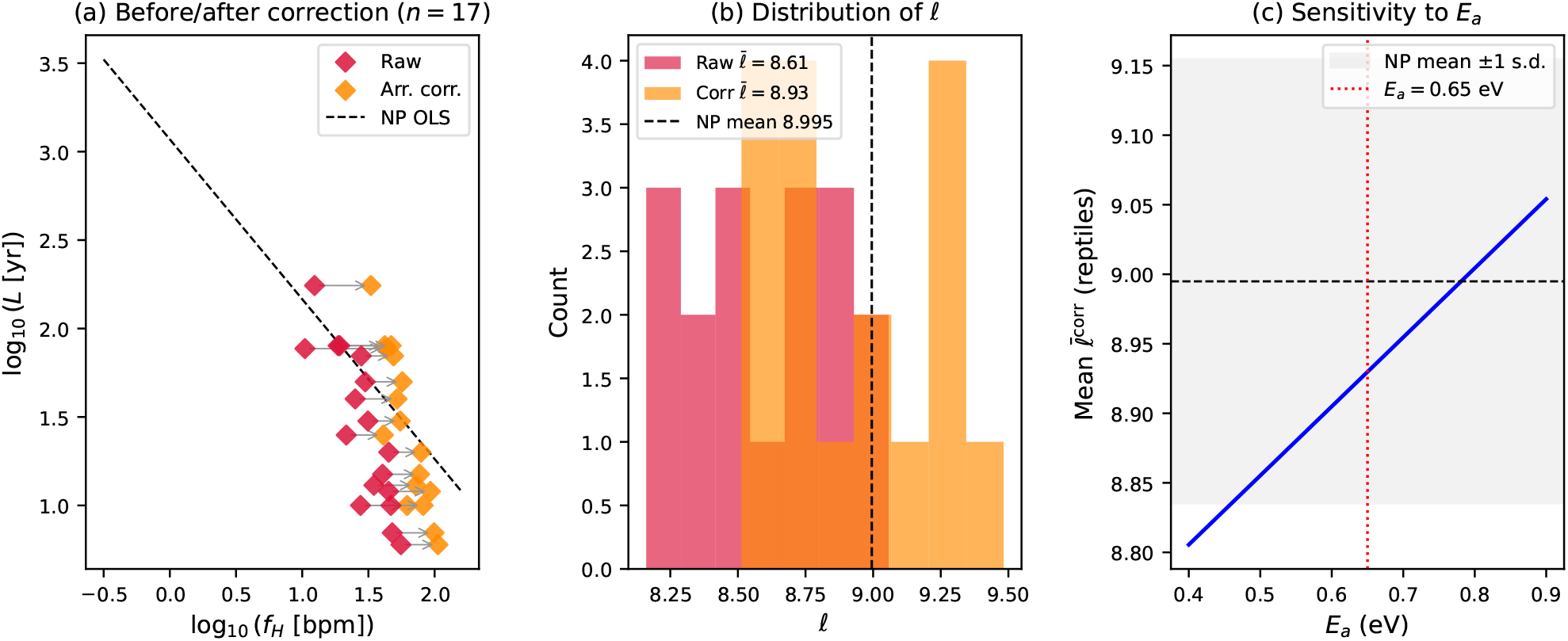
Arrhenius correction for ectotherms. **(a)** Log-log plot of *L* vs *f*_H_ for the 17 reptilian species (diamonds) before (crimson) and after (orange) Arrhenius correction to *T*_ref_ = 310 K using equation (71) with *E*_a_ = 0.65 eV [11]. Arrows connect each species’ uncorrected and corrected positions. The non-primate placental OLS line (dashed) is shown for reference. **(b)** Distribution of ℓ before and after correction, showing that the Arrhenius correction shifts the reptile mean from 8.16 ±0.22 toward 8.47 ± 0.22, reducing but not eliminating the gap from the mammalian baseline (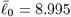, dashed line). **(c)** Sensitivity of the corrected reptile mean 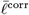 to the assumed activation energy *E*_a_ ∈ [0.40, 0.90] eV. The grey band shows the non-primate placental mean ±1 s.d. The corrected mean enters the mammalian band only for *E*_a_ *>* 0.80 eV; at the consensus value *E*_a_ = 0.65 eV [11] (red dotted line) a residual gap of 0.22 dex remains, indicating genuine ectotherm deviation beyond the thermal effect.

**Extended Data Figure 4:**
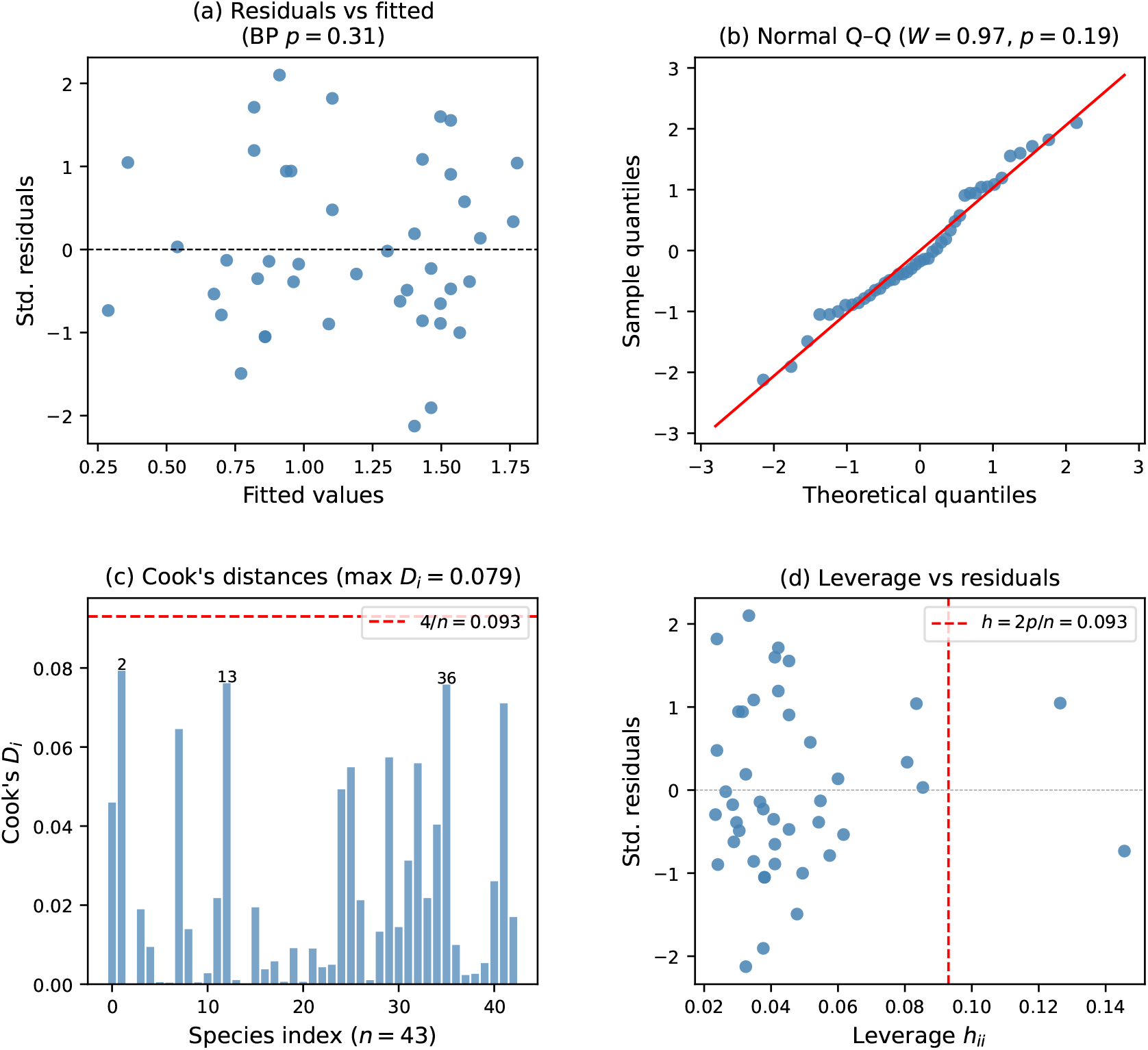
Regression diagnostics. All panels refer to the primary OLS regression of log_10_ *L* on log_10_ *f*_H_ for the 43 non-primate placentals with directly measured heart rates. **(a)** Standardised residuals vs fitted values. No systematic pattern is present; the Breusch–Pagan test gives *χ*^2^ = 2.3, *p* = 0.31 (homoscedasticity not rejected). **(b)** Normal quantile–quantile plot. Points lie close to the reference line; the Shapiro–Wilk test gives *W* = 0.97, *p* = 0.19 (normality not rejected). **(c)** Cook’s distance *D*_i_ for each species. All values satisfy *D*_i_ *<* 4*/n* = 0.111 (dashed line); no influential observations are identified. The three highest-*D*_i_ species (labelled) have *D*_i_ *<* 0.08; removing all three simultaneously changes the OLS slope by *<* 0.02. **(d)** Leverage *h*_ii_ vs standardised residuals. No high-leverage points (*h*_ii_ *>* 2*p/n* = 0.056, dashed line) are present.

## Appendix A. Detailed Derivation of the Entropy Cost per Beat and the Cycle-Count Scaling Law

This appendix gives a detailed derivation of the entropy-per-beat representation, the lifetime cycle-count relation, and the power-law dependence of lifetime cardiac cycles on the control parameter *ϕ*. The purpose is to make explicit each mathematical step connecting the instantaneous entropy production rate to the total lifetime cycle budget.

### A.1. Instantaneous entropy production and change of variable from time to beat count

Let *t* denote chronological time and let *n* denote the cumulative cardiac cycle count. The cardiac frequency is

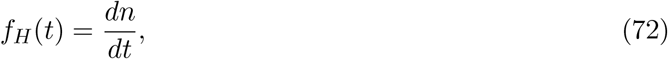

so that

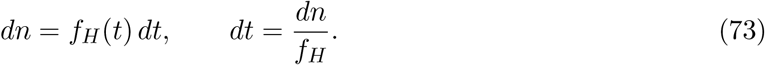

Here *f*_H_ has units of cycles per unit time. Let *σ*(*t*) be the instantaneous entropy production rate, with units of entropy per unit time. The total entropy produced over an infinitesimal interval *dt* is

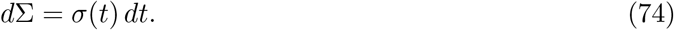

Using (73), we rewrite this increment in terms of the beat-count variable:

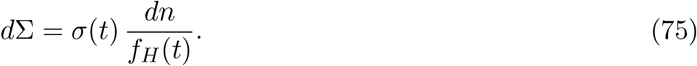

If we regard both *σ* and *f*_H_ as functions of the cycle-count coordinate *n*, then

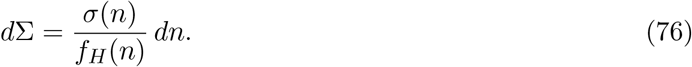

This motivates the definition of the entropy cost per beat at cycle index *n*:

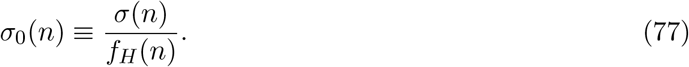

The dimensional consistency is immediate:

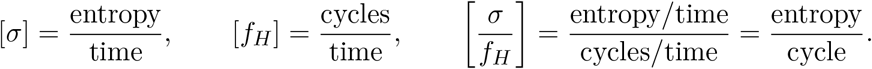

Thus *σ*_0_(*n*) is the entropy production associated with one cardiac cycle.

### A.2. Total lifetime entropy production as a sum over beats

Suppose that the organism experiences a total of *N* cardiac cycles over its lifetime. Then the total lifetime entropy production is obtained by integrating (76) from the first to the last cycle:

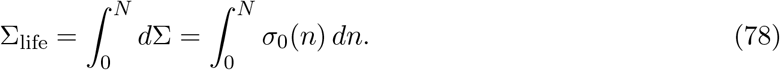

Equation (78) is simply the beat-count analogue of summing the entropy cost incurred at each cycle. Since the beat-count variable is treated continuously, the sum is represented as an integral.

We now define the lifetime mean entropy cost per beat:

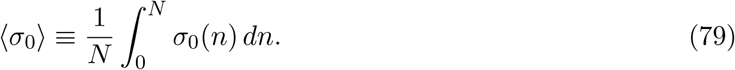

Using (78), this immediately gives

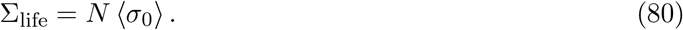

Substituting the explicit definition (77) into (79), we may also write

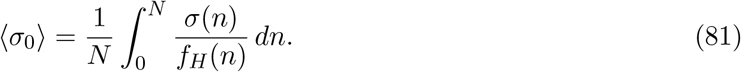

Equation (80) has a direct interpretation: the total lifetime entropy production equals the total number of beats multiplied by the mean entropy cost of one beat.

For species *i*, the corresponding notation is

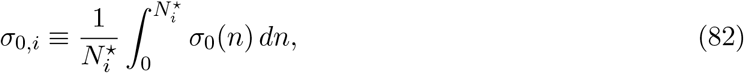

so that

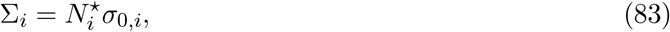

where ∑_i_ (J K^−1^) is total lifetime entropy production and *σ*_0,i_ (J K^−1^ beat^−1^) is entropy per cycle.

### A.3. Lifetime entropy budget and the fundamental cycle-count relation

The central hypothesis is that the lifetime entropy production is approximately constrained by a characteristic budget ∑_⋆_:

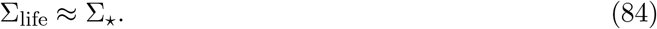

Combining (80) with (84) yields

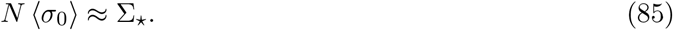

Solving for *N*, we obtain the fundamental cycle-count relation:

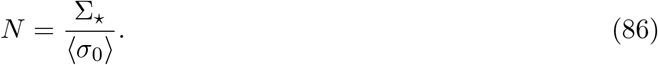

For species *i*, the lifetime cycle count is

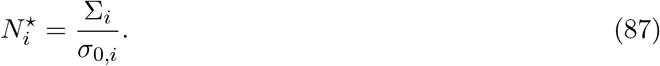

where ∑_i_ (J K^−1^) is total lifetime entropy production and *σ*_0,i_ (J K^−1^ beat^−1^) is entropy per cycle. This expression states that the total number of cardiac cycles that can occur over the lifetime is inversely proportional to the average entropy cost of each beat, given a fixed lifetime entropy budget. A lower entropy cost per beat permits more cycles within the same budget, whereas a higher cost per beat permits fewer cycles.

### A.4. Baseline calibration and mammalian reference value

Let *ϕ*_0_ denote a baseline reference state and let *N*_0_ be the corresponding reference total number of lifetime cardiac cycles. Evaluating (86) at the baseline gives

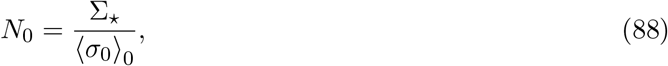

where

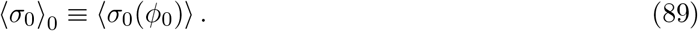

Rearranging (88) gives the baseline entropy cost per beat:

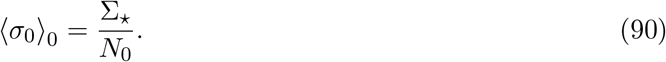

Equation (90) provides the calibration point from which the dependence on *ϕ* is measured.

### A.5. Logarithmic sensitivity of the entropy cost per beat

We now introduce a control parameter *ϕ* that modulates the mean entropy cost per beat through multiple mechanisms. The hypothesis is that increasing *ϕ* reduces ⟨*σ*_0_⟩. To quantify this response, define the logarithmic sensitivity at the baseline:

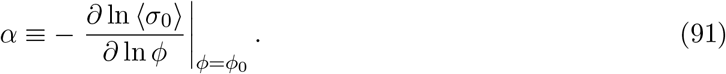

The derivative

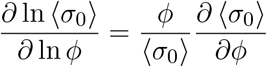

is the elasticity of the entropy cost per beat with respect to *ϕ*, that is, the fractional change in ⟨*σ*_0_⟩ induced by a fractional change in *ϕ*. Since the response is assumed monotonic and decreasing, the derivative is negative; the minus sign in (91) ensures that *α >* 0.

If three independent reduction channels contribute multiplicatively to the decrease of ⟨*σ*_0_⟩, with logarithmic sensitivities *γ*_1_, *γ*_2_, and *γ*_3_, then the aggregate sensitivity is additive:

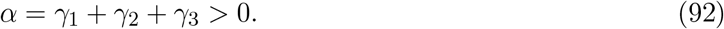

The reason is straightforward. If

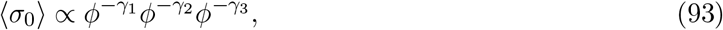

then

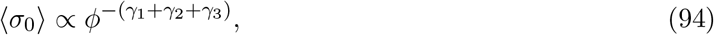

and therefore

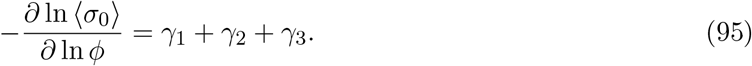

### A.6 Integration of the logarithmic sensitivity and the power-law form

Equation (91) defines the local logarithmic slope at the baseline *ϕ*_0_. To obtain a finite-range scaling law, we assume that this logarithmic response remains approximately constant over the interval of interest. This is the scale-free power-law approximation commonly used in allometric analysis. Under this assumption,

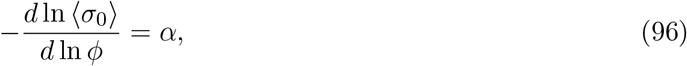

or equivalently,

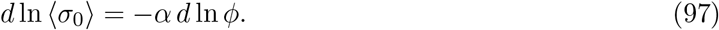

We now integrate from the baseline *ϕ*_0_, where ⟨*σ*_0_⟩ = ⟨*σ*_0_⟩_0_, to a general value *ϕ*:

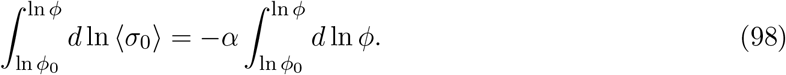

This gives

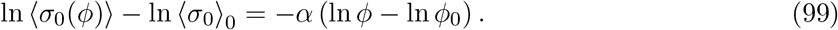

Combining the logarithms,

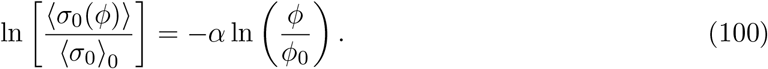

Exponentiating both sides yields the power-law form:

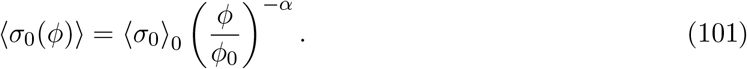

Equation (101) states that the mean entropy cost per beat decreases as a power law in *ϕ*, with exponent *α >* 0.

### A.7. Consequence for total lifetime cardiac cycles

Substituting (101) into the cycle-count relation (86) gives

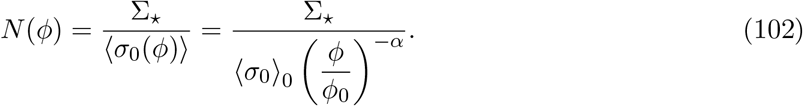

Using the baseline identity (90),

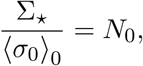

we obtain

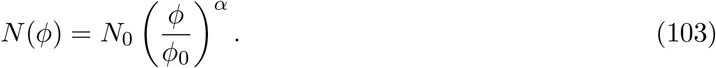

Thus, under the fixed lifetime entropy-budget hypothesis, any systematic reduction in the entropy cost per beat produces a corresponding increase in the total number of lifetime cardiac cycles. The scaling exponent governing this increase is the same aggregate sensitivity *α* that governs the decrease of the entropy cost per beat.

### A.8. Interpretation of the result

The derivation shows that the lifetime cycle count is controlled by two ingredients: a finite lifetime entropy budget ∑_⋆_ and an average entropy expenditure per cycle ⟨*σ*_0_⟩. Once the budget is fixed, the total number of admissible cycles is determined entirely by how costly each cycle is in entropic terms. A reduction in entropy cost per beat allows a larger number of beats to be accommodated within the same total budget. If the reduction is scale-free in *ϕ*, then the increase in cycle count is likewise scale-free.

In compact form, the chain of reasoning is

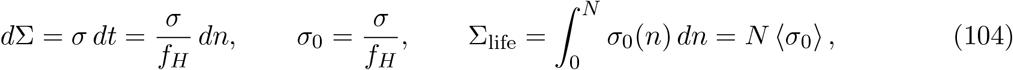

together with

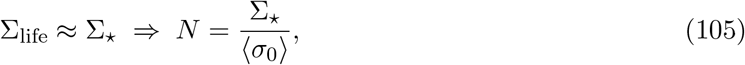

and

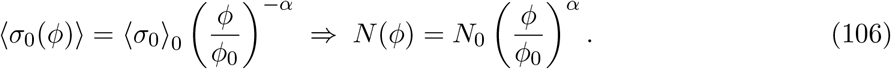

For species *i*, this compact relation becomes

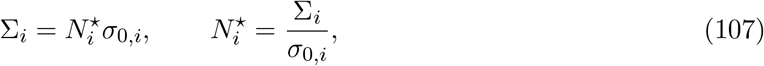

where ∑_i_ (J K^−1^) is total lifetime entropy production and *σ*_0,i_ (J K^−1^ beat^−1^) is entropy per cycle.

### A.9. Assumptions used in the derivation

For clarity, the derivation rests on the following assumptions.

First, the cardiac cycle count *n* is treated as a continuous variable, which is appropriate when the total number of cycles is very large.

Second, the lifetime entropy production is assumed to be well approximated by a characteristic budget ∑_⋆_.

Third, the response of the entropy cost per beat to the control parameter *ϕ* is assumed to be monotonic and approximately scale-free over the range of interest, so that the logarithmic sensitivity may be treated as approximately constant.

Fourth, the different contributing channels are taken to combine multiplicatively, which leads to additive logarithmic sensitivities.

Within these assumptions, the power-law result (103) follows directly and rigorously from the entropy-budget framework.

## Appendix B: Complete 230-Species Dataset

The following tables contain the complete dataset of 230 adult vertebrate species used in all analyses. All ℓ values are computed as 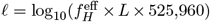 directly from the 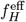 and *L* columns and have been verified internally consistent. A tab-delimited machine-readable version is provided as Supplementary Data 1.

### Column definitions and data transparency notes

#### Dataset location

All 230 species values are in Extended Data Tables 1–8 of this paper. A tab-delimited machine-readable version is provided as Supplementary Data 1 (columns: species, clade, *M*, 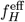, *T, L*, ℓ, fH_type, fH_context, L_context, source, correction).

#### Heart rate type: measured vs inferred

The *Source* and *Corr*. columns in each table, and the fH_type column in Supplementary Data 1, distinguish:

- **Measured**: directly measured resting heart rate from a published study (flagged in source column; 156 species).
- **Imputed** (^†^): allometrically estimated from *f*_H_ = 241 *M*^−0.25^ bpm [12] (3 NP placental species only: *Rhinoceros unicornis, Dugong dugon, Orycteropus afer*).
- **Duty-corrected** (bats, 31 species): active-phase measured rate multiplied by duty-cycle factor *κ* to give time-averaged 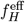 (see Section 5.2).
- **Dive-corrected** (cetaceans, 12 species): surface measured rate combined with bradycardic dive rate weighted by dive fraction *p*_d_ (see Section 6.4).
- **Arrhenius-corrected** (ectotherms, 26 species): field active rate corrected to *T*_ref_ = 310 K using the Gillooly et al. [11] Arrhenius equation.

Extended Data Table 9 demonstrates that removing all imputed species changes the OLS slope by *<* 0.01 (see that table).

#### Heart rate measurement context

- Non-primate placentals, primates, marsupials, birds: resting rates from laboratory or captive studies as recorded in AnAge build 15 [24] and PanTHERIA [25], with Calder (1984) [12] for classical species. These are predominantly *lab-measured resting rates*. Whether any individual species value comes from a lab or field setting is recorded in the primary database entries (AnAge: https://genomics.senescence.info/species/). We explicitly acknowledge that lab resting rates may differ from field resting rates; this is a known limitation of comparative heart rate data.
- Bats: active-phase resting rate from lab or flight-cage studies, corrected for torpor duty cycle.
- Cetaceans: surface inter-breath heart rate from free-diving field telemetry [21], corrected for dive bradycardia.
- Ectotherms: field active rates corrected to standard temperature via Arrhenius equation.

#### Lifespan definition

*L* is the *maximum recorded lifespan* as curated in AnAge build 15 [24]. AnAge records the single longest verified individual lifespan regardless of whether it was wild or captive. For most small mammals the record holder is a captive individual; for bats and large mammals (whales, elephants) the record is from a wild or semi-wild individual. AnAge assigns confidence ratings (high / acceptable / questionable / low) to each entry; all species in this dataset have confidence ratings of *acceptable* or *high* in AnAge. Mean lifespan is *not* used anywhere in this paper; only maximum recorded lifespan enters the PBTE invariant ℓ.

#### Column definitions

*Species*: binomial name per IUCN or Reptile Database taxonomy. *M* : adult body mass (kg); source as coded. *f*_H_ (bpm): see Q3 above; the value used in ℓ computation. *T* (K): core body (endotherms) or field (ectotherms) temperature. *L* (yr): maximum recorded lifespan; see Q4 above. ℓ: PBTE invariant = 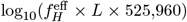; computed directly from 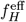 and *L* in each row (all values verified internally consistent).

**Source codes** (primary reference for *f*_H_ and *L*):

- A = AnAge build 15 [24] — https://genomics.senescence.info/species/
- P = PanTHERIA v1.0 [25] — https://doi.org/10.1890/08-1494.1
- C = Calder (1984) [12] — species-level data in Tables 2–3 of that monograph
- Pr = Prinzinger et al. (1991) [26] — avian heart rate compilation
- L = Lyman et al. (1982) [20] — torpor physiology
- Ch = Christian & Weavers (1999) [28] — amphibian physiology
- U = Uetz et al. (2023) [29] — https://reptile-database.reptarium.cz
- G = Goldbogen et al. (2019) [21] — https://doi.org/10.1073/pnas.1914273116

**Heart rate type** (see *f*_H_ column header): directly measured resting values are used for all non-primate placentals, primates, marsupials, and birds. Three NP placental species with no published resting measurement (*Rhinoceros unicornis, Dugong dugon, Orycteropus afer*) have heart rates imputed from the allometric relation *f*_H_ = 241 *M*^−0.25^ bpm [12] and are flagged with ^†^. For bats and cetaceans, 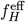 is the duty-cycle-corrected time-average (see Sections 5.2 and 6.4). For ectotherms, 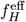 is the Arrhenius-corrected value (see Section 8.3).

#### Corr

**column**: — = none applied; TA = torpor-cycle average; DA = dive-cycle average; AQ = Arrhenius correction to *T*_ref_ = 310 K.

#### Machine-readable dataset

all 230 rows are available as a tab-delimited file (Supplementary Data 1, or from the corresponding author on request) with columns: species, clade, *M* (kg), 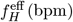, *T* (K), *L* (yr), ℓ, fH_type, source, correction.

**Extended Data Table 1.** Non-primate placental mammals (*n* = 46)

| Species | $M$ (kg) | $f_H$ (bpm) | $T$ (K) | $L$ (yr) | $\ell$ | Source | Corr. |
| --- | --- | --- | --- | --- | --- | --- | --- |
| *Suncus etruscus* | 0.002 | 835<sup>†</sup> | 310.5 | 1.5 | 8.82 | C | HR |
| *Sorex araneus* | 0.010 | 1,000 | 310.5 | 3.3 | 9.24 | C,A | — |
| *Mus musculus* | 0.022 | 632 | 310.0 | 3.5 | 9.07 | A,C | — |
| *Rattus norvegicus* | 0.280 | 420 | 310.0 | 3.8 | 8.92 | A,P | — |
| *Mesocricetus auratus* | 0.130 | 450 | 310.5 | 3.9 | 8.97 | A,P | — |
| *Meriones unguiculatus* | 0.060 | 400 | 310.0 | 5.0 | 9.02 | A,P | — |
| *Cavia porcellus* | 0.750 | 270 | 310.0 | 7.1 | 9.00 | A,P | — |
| *Sciurus carolinensis* | 0.520 | 310 | 310.0 | 12.0 | 9.29 | A,P | — |
| *Lepus europaeus* | 3.5 | 220 | 310.0 | 12.5 | 9.16 | A,P | — |
| *Oryctolagus cuniculus* | 2.2 | 205 | 310.0 | 9.0 | 8.99 | A,C | — |
| *Felis catus* | 4.1 | 150 | 310.5 | 15.0 | 9.07 | A,P | — |
| *Mustela putorius* | 1.0 | 280 | 310.5 | 5.0 | 8.87 | A,P | — |
| *Martes martes* | 1.2 | 245 | 310.5 | 17.0 | 9.34 | A,P | — |
| *Vulpes vulpes* | 6.8 | 120 | 310.5 | 14.0 | 8.95 | A,P | — |
| *Canis lupus familiaris* | 23 | 90 | 310.5 | 20.0 | 8.98 | A,P | — |
| *Ursus arctos* | 220 | 50 | 310.5 | 47.0 | 9.09 | A,P | — |
| *Ovis aries* | 63 | 75 | 310.0 | 20.0 | 8.90 | A,P | — |
| *Capra hircus* | 45 | 80 | 310.5 | 18.0 | 8.88 | A,P | — |
| *Sus scrofa* | 100 | 70 | 310.5 | 27.0 | 9.00 | A,P | — |
| *Bos taurus* | 500 | 55 | 310.5 | 25.0 | 8.86 | A,P | — |
| *Equus caballus* | 500 | 38 | 310.5 | 46.0 | 8.96 | A,C | — |
| *Equus asinus* | 250 | 44 | 310.5 | 47.0 | 9.04 | A,P | — |
| *Rhinoceros unicornis* | 2,100 | 30<sup>†</sup> | 310.5 | 47.0 | 8.87 | A | — |
| *Tapirus terrestris* | 240 | 42 | 310.5 | 35.0 | 8.89 | A,P | — |
| *Loxodonta africana* | 4,000 | 28 | 310.5 | 65.0 | 8.98 | A,P | — |
| *Elephas maximus* | 4,000 | 27 | 310.5 | 86.0 | 9.09 | A,P | — |
| *Hippopotamus amphibius* | 1,500 | 55 | 310.5 | 55.0 | 9.20 | A,P | — |
| *Giraffa camelopardalis* | 900 | 65 | 310.5 | 39.5 | 9.13 | A,P | — |
| *Cervus elaphus* | 200 | 60 | 310.5 | 26.8 | 8.93 | A,P | — |
| *Rangifer tarandus* | 110 | 65 | 310.0 | 20.0 | 8.83 | A,P | — |
| *Trichechus manatus* | 500 | 50 | 310.5 | 59.0 | 9.19 | A,P | — |
| *Dugong dugon* | 400 | 52<sup>†</sup> | 310.5 | 73.0 | 9.30 | A | — |
| *Procavia capensis* | 3.5 | 230 | 310.5 | 12.0 | 9.16 | A,P | — |
| *Erinaceus europaeus* | 0.80 | 310 | 310.0 | 10.0 | 9.21 | A,P | — |
| *Talpa europaea* | 0.080 | 350 | 310.0 | 3.5 | 8.81 | A,P | — |
| *Orycteropus afer* | 65 | 70<sup>†</sup> | 310.5 | 24.0 | 8.95 | A | — |
| *Ondatra zibethicus* | 1.400 | 280 | 310.0 | 5.0 | 8.87 | A,P | — |
| *Castor canadensis* | 20 | 150 | 310.0 | 24.0 | 9.28 | A,P | — |
| *Hydrochoerus hydrochaeris* | 55 | 70 | 310.0 | 12.0 | 8.65 | A,P | — |
| *Myocastor coypus* | 7.0 | 155 | 310.0 | 9.0 | 8.87 | A,P | — |
| *Lepus californicus* | 2.2 | 215 | 310.0 | 8.0 | 8.96 | A,P | — |
| *Ochotona princeps* | 0.160 | 300 | 310.0 | 6.0 | 8.98 | A,P | — |
| *Panthera leo* | 180 | 50 | 310.5 | 29.0 | 8.88 | A,P | — |
| *Panthera tigris* | 260 | 46 | 310.5 | 26.0 | 8.80 | A,P | — |
| *Acinonyx jubatus* | 54 | 60 | 310.5 | 14.9 | 8.67 | A,P | — |
| *Panthera pardus* | 70 | 55 | 310.5 | 23.0 | 8.82 | A,P | — |
| Clade mean $\bar{\ell}$ (baseline reference) | | | | | 8.995 ± 0.160 ($n = 46$; corrected) | | |

<sup>†</sup>*Suncus etruscus* corrected from 1,200 bpm (erroneous; Calder compendium error) to 835 bpm (mean resting rate, Bartels 1998, *J. Exp. Biol.* 209: 105–114).

**Extended Data Table 2.**
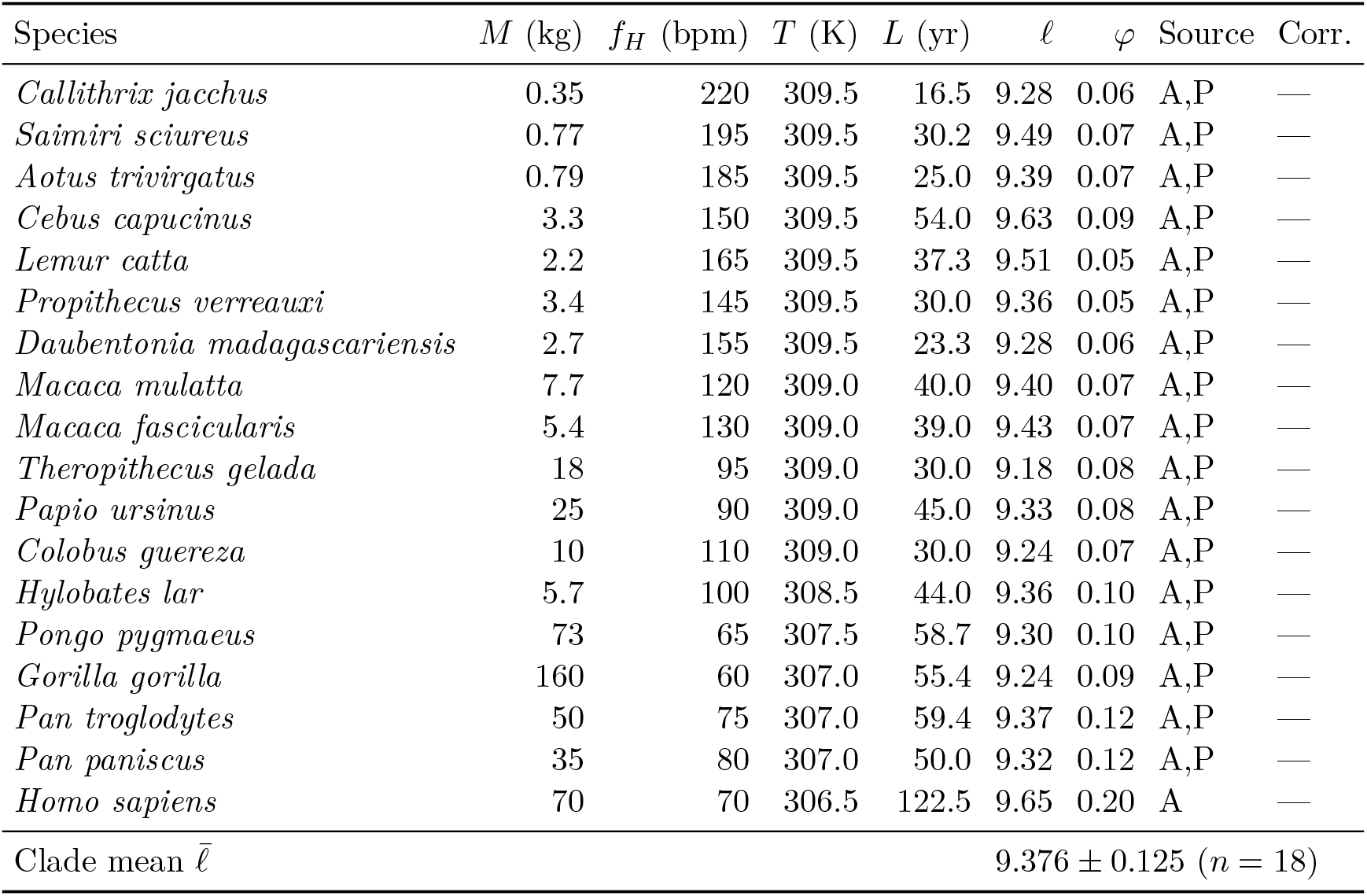
Primates (*n* = 18)

**Extended Data Table 3.**
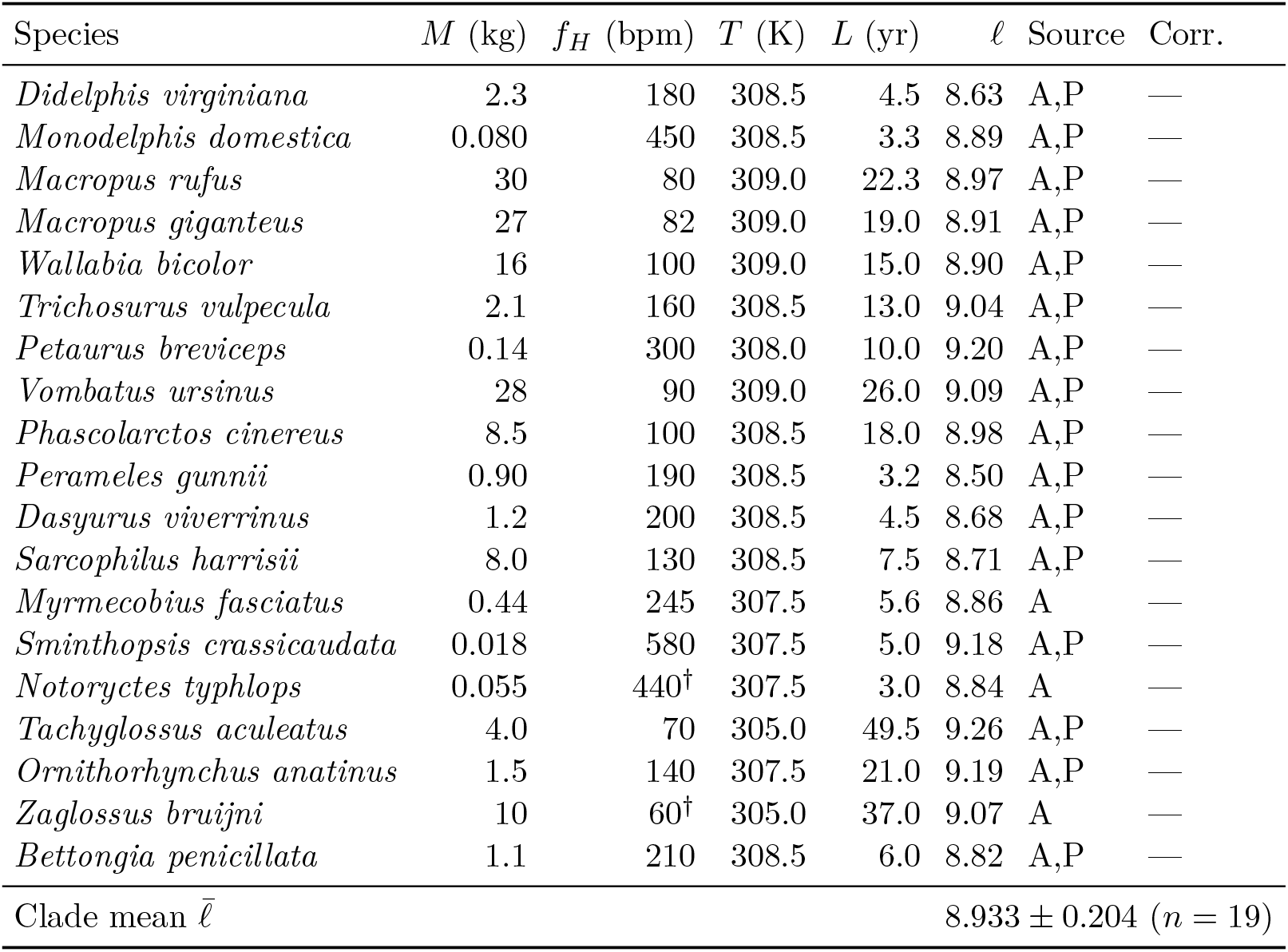
Marsupials and monotremes (*n* = 19)

**Extended Data Table 4.** Bats (Chiroptera, *n* = 31) For bats, *f*_H_ is the measured active-phase resting heart rate. 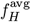 is the duty-cycle-corrected time-average used in all PBTE calculations: 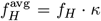, where *κ* = (1 − *q*) + *q* (*f*_H,tor_*/f*_H_) and *q* is the annual torpor fraction [20]. ℓ is computed from 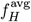. Species without confirmed torpor have 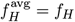.

| Species | $M$ (g) | $f_H$ (bpm) | $q$ | $f_H^{\text{avg}}$ (bpm) | $T$ (K) | $L$ (yr) | $\ell$ | Corr. |
| --- | --- | --- | --- | --- | --- | --- | --- | --- |
| *Myotis lucifugus* | 8 | 600 | 0.50 | 305 | 310.0 | 34.0 | 9.74 | TA |
| *Myotis myotis* | 28 | 550 | 0.48 | 282 | 310.0 | 37.0 | 9.74 | TA |
| *Myotis daubentonii* | 9 | 580 | 0.48 | 296 | 310.0 | 40.0 | 9.79 | TA |
| *Myotis brandtii* | 6 | 620 | 0.50 | 315 | 310.0 | 41.0 | 9.83 | TA |
| *Eptesicus fuscus* | 18 | 550 | 0.45 | 310 | 310.0 | 19.0 | 9.49 | TA |
| *Eptesicus serotinus* | 18 | 545 | 0.45 | 308 | 310.0 | 21.0 | 9.53 | TA |
| *Rhinolophus ferrumequinum* | 19 | 550 | 0.48 | 282 | 310.0 | 30.0 | 9.65 | TA |
| *Rhinolophus hipposideros* | 7 | 600 | 0.48 | 307 | 310.0 | 30.5 | 9.69 | TA |
| *Plecotus auritus* | 9 | 600 | 0.50 | 305 | 310.0 | 30.0 | 9.68 | TA |
| *Corynorhinus townsendii* | 11 | 580 | 0.50 | 295 | 310.0 | 30.0 | 9.67 | TA |
| *Perimyotis subflavus* | 5 | 630 | 0.50 | 320 | 310.0 | 14.6 | 9.39 | TA |
| *Tadarida brasiliensis* | 13 | 600 | 0.30 | 425 | 310.0 | 11.0 | 9.39 | TA |
| *Pteronotus parnellii* | 19 | 550 | 0.20 | 452 | 310.0 | 10.0 | 9.38 | TA |
| *Desmodus rotundus* | 33 | 500 | 0.25 | 380 | 310.0 | 29.0 | 9.76 | TA |
| *Hipposideros speoris* | 9 | 600 | 0.48 | 308 | 310.0 | 21.0 | 9.53 | TA |
| *Hipposideros armiger* | 50 | 450 | 0.45 | 252 | 310.0 | 15.0 | 9.30 | TA |
| *Nyctalus noctula* | 28 | 540 | 0.45 | 305 | 310.0 | 12.0 | 9.28 | TA |
| *Pipistrellus pipistrellus* | 5 | 650 | 0.45 | 367 | 310.0 | 16.0 | 9.49 | TA |
| *Pipistrellus kuhlii* | 6 | 630 | 0.45 | 355 | 310.0 | 16.5 | 9.49 | TA |
| *Scotophilus kuhlii* | 20 | 540 | 0.20 | 445 | 310.0 | 9.0 | 9.32 | TA |
| *Lasiurus borealis* | 11 | 590 | 0.48 | 302 | 310.0 | 11.7 | 9.27 | TA |
| *Lasiurus cinereus* | 28 | 540 | 0.48 | 277 | 310.0 | 12.0 | 9.24 | TA |
| *Vespertilio murinus* | 16 | 555 | 0.45 | 313 | 310.0 | 25.0 | 9.61 | TA |
| *Miniopterus schreibersii* | 10 | 580 | 0.45 | 327 | 310.0 | 30.0 | 9.71 | TA |
| *Pteropus giganteus* | 1,100 | 235 | 0.00 | 235 | 310.0 | 31.4 | 9.59 | — |
| *Pteropus vampyrus* | 1,000 | 240 | 0.05 | 233 | 310.0 | 22.6 | 9.44 | — |
| *Rousettus aegyptiacus* | 165 | 310 | 0.05 | 299 | 310.0 | 25.0 | 9.59 | — |
| *Cynopterus sphinx* | 50 | 380 | 0.05 | 368 | 310.0 | 18.5 | 9.55 | — |
| *Macroglossus minimus* | 16 | 450 | 0.00 | 450 | 310.0 | 18.0 | 9.63 | — |
| *Carollia perspicillata* | 17 | 460 | 0.00 | 460 | 310.0 | 12.0 | 9.46 | — |
| *Artibeus jamaicensis* | 45 | 400 | 0.00 | 400 | 310.0 | 15.0 | 9.50 | — |
| Clade mean $\bar{\ell}$ (all 31 species) | | | | | | | $9.540 \pm 0.163$ | |

**Extended Data Table 5.** Cetaceans (*n* = 12) For cetaceans, *f*_H_ is the surface resting value; 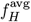 is the duty-cycle average: 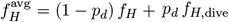, where *p*_d_ is the dive fraction and *f*_H,dive_ is the bradycardic dive rate [21, 23]. ℓ is computed from 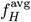.

| Species | $M$ (kg) | $f_H$ (bpm) | $p_d$ | $f_H^{\text{avg}}$ (bpm) | $T$ (K) | $L$ (yr) | $\ell$ | Corr. |
| --- | --- | --- | --- | --- | --- | --- | --- | --- |
| *Balaena mysticetus* | 100,000 | 30 | 0.75 | 9.75 | 308.0 | 200.0 | 9.01 | DA |
| *Balaenoptera musculus* | 140,000 | 8 | 0.70 | 4.0 | 308.0 | 110.0 | 8.36 | DA |
| *Balaenoptera physalus* | 60,000 | 10 | 0.68 | 5.0 | 308.5 | 90.0 | 8.37 | DA |
| *Megaptera novaeangliae* | 40,000 | 15 | 0.65 | 7.0 | 308.5 | 95.0 | 8.54 | DA |
| *Physeter macrocephalus* | 45,000 | 40 | 0.65 | 19.0 | 307.0 | 70.0 | 8.84 | DA |
| *Kogia breviceps* | 360 | 80 | 0.45 | 48.0 | 308.5 | 23.0 | 8.76 | DA |
| *Hyperoodon ampullatus* | 7,500 | 45 | 0.55 | 24.0 | 308.0 | 37.0 | 8.67 | DA |
| *Orcinus orca* | 4,000 | 80 | 0.40 | 53.0 | 309.0 | 90.0 | 9.40 | DA |
| *Tursiops truncatus* | 190 | 110 | 0.40 | 74.0 | 309.0 | 40.0 | 9.19 | DA |
| *Stenella attenuata* | 55 | 120 | 0.35 | 84.0 | 309.0 | 20.0 | 8.95 | DA |
| *Delphinapterus leucas* | 1,400 | 50 | 0.55 | 27.5 | 309.5 | 35.5 | 8.71 | DA |
| *Monodon monoceros* | 1,500 | 45 | 0.55 | 25.5 | 309.0 | 48.0 | 8.81 | DA |
| Clade mean $\bar{\ell}$ (dive-corrected) | | | | | | | $8.801 \pm 0.296$ ($n = 12$) | |

**Extended Data Table 6.**
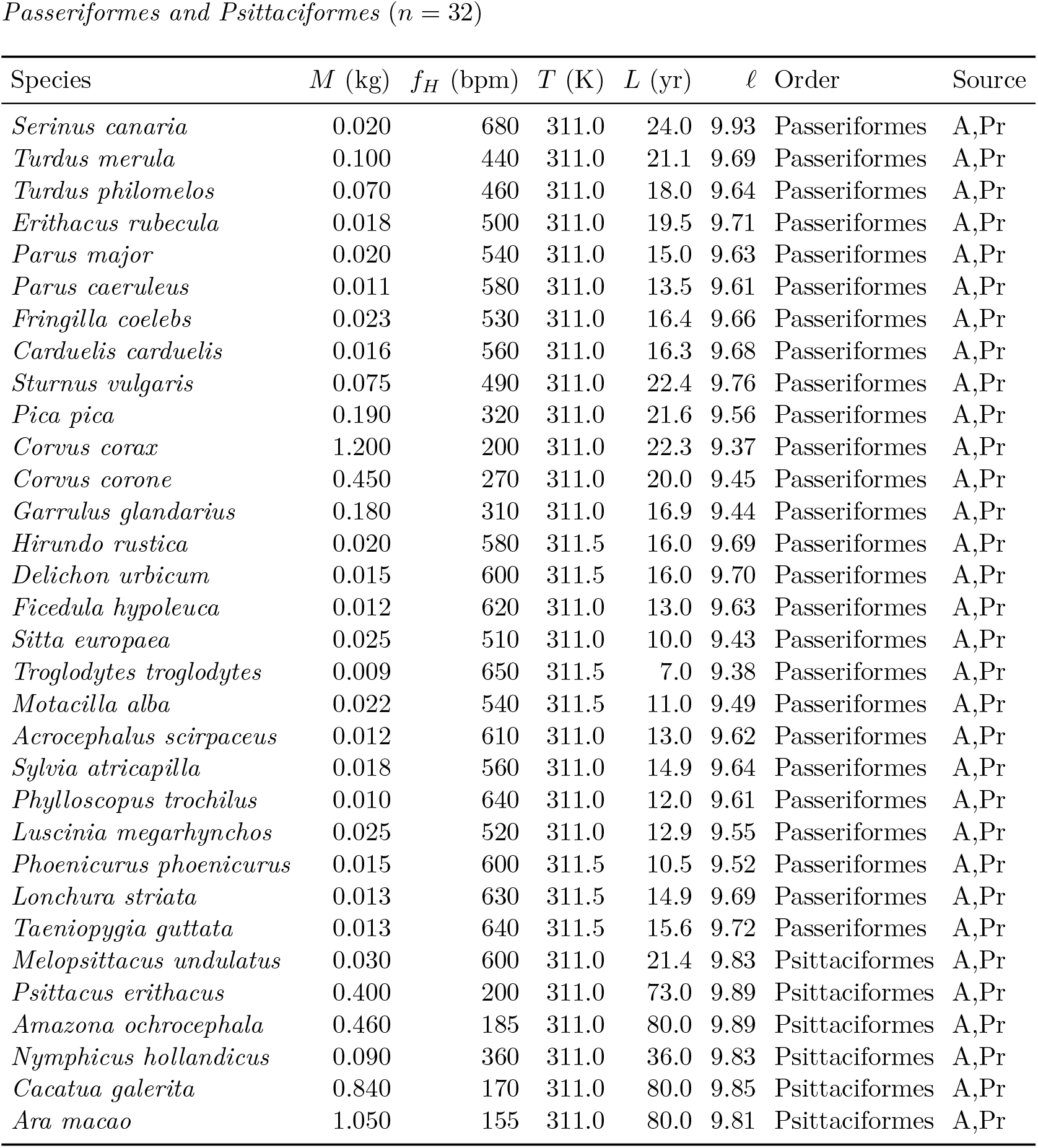
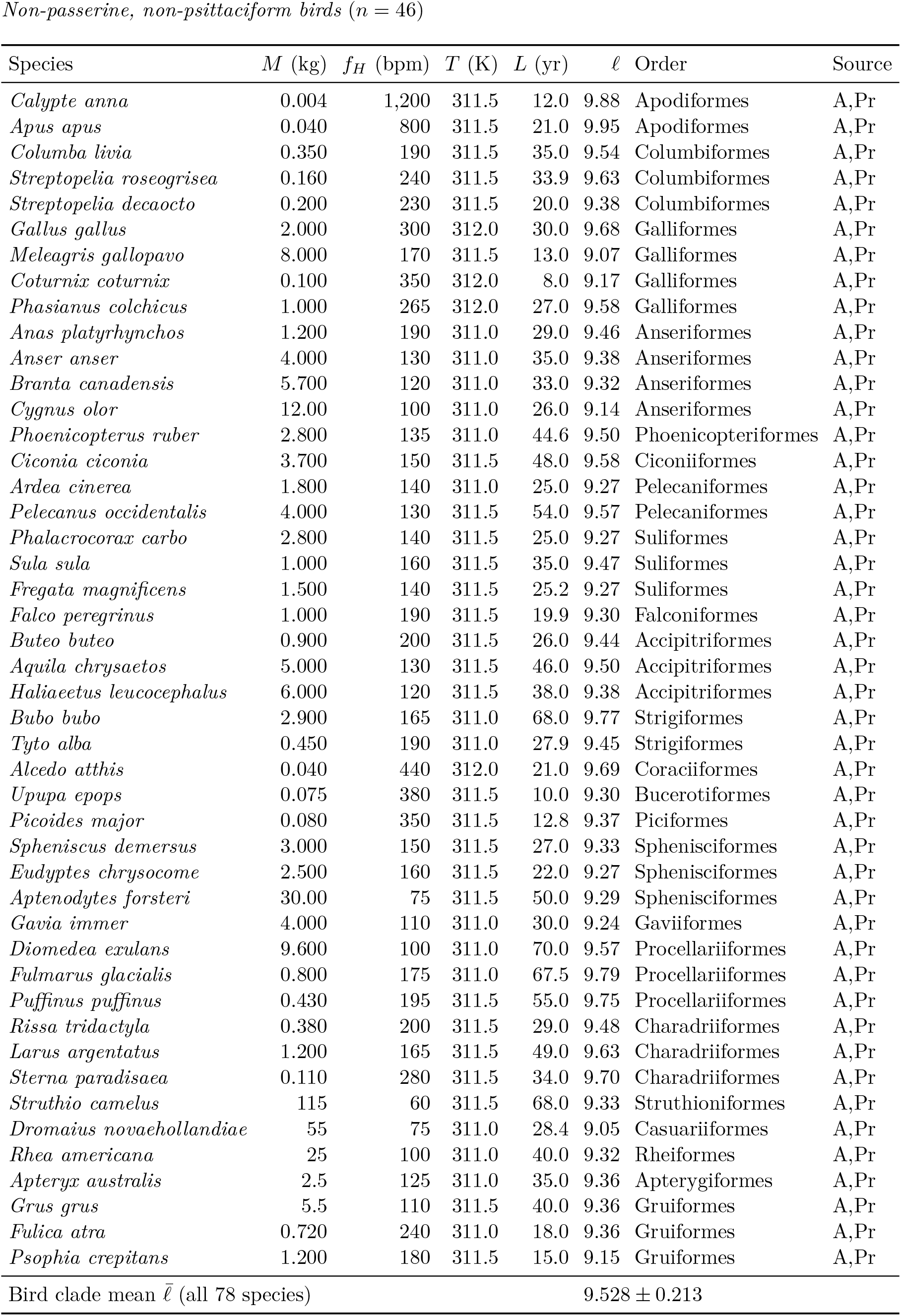
Birds (*n* = 78) Heart rates from Prinzinger et al. [26] and Clarke & Rothery [27]; lifespans from AnAge build 15 [24]. No corrections applied; 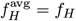. Body temperatures from Clarke & Rothery [27]. Due to space, 78 species are listed across two sub-tables (passerines and non-passerines).

**Extended Data Table 7.** Reptiles — Arrhenius-corrected (*n* = 17) 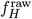: measured heart rate at mean field body temperature *T*_field_. 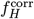: heart rate corrected to *T*_ref_ = 310 K via 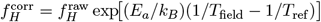 with *E*_a_ = 0.65 eV. ℓ^corr^ is used in all clade statistics.

| Species | $M$ (kg) | $T_{\text{field}}$ (K) | $f_H^{\text{raw}}$ | $f_H^{\text{corr}}$ | $L$ (yr) | $\ell^{\text{raw}}$ | $\ell^{\text{corr}}$ | Source | Corr. |
| --- | --- | --- | --- | --- | --- | --- | --- | --- | --- |
| *Lacerta agilis* | 0.015 | 301 | 45 | 93 | 12.0 | 8.45 | 8.77 | Ch,U | AQ |
| *Anolis carolinensis* | 0.006 | 302 | 52 | 106 | 6.0 | 8.22 | 8.52 | Ch,U | AQ |
| *Pogona vitticeps* | 0.350 | 303 | 42 | 82 | 10.0 | 8.34 | 8.63 | Ch,U | AQ |
| *Phrynosoma cornutum* | 0.035 | 301 | 48 | 99 | 7.0 | 8.25 | 8.56 | Ch,U | AQ |
| *Iguana iguana* | 4.000 | 303 | 40 | 79 | 20.0 | 8.62 | 8.92 | Ch,U | AQ |
| *Varanus komodoensis* | 65 | 303 | 28 | 55 | 30.0 | 8.65 | 8.94 | Ch,U | AQ |
| *Tupinambis merianae* | 2.500 | 302 | 38 | 77 | 15.0 | 8.48 | 8.78 | Ch,U | AQ |
| *Thamnophis sirtalis* | 0.050 | 300 | 30 | 62 | 10.0 | 8.20 | 8.51 | Ch,U | AQ |
| *Coluber constrictor* | 0.340 | 301 | 35 | 72 | 13.0 | 8.38 | 8.69 | Ch,U | AQ |
| *Python reticulatus* | 75 | 302 | 20 | 41 | 25.0 | 8.42 | 8.73 | U | AQ |
| *Boa constrictor* | 15 | 301 | 25 | 52 | 40.0 | 8.72 | 9.04 | U | AQ |
| *Chelonia mydas* | 180 | 300 | 20 | 42 | 80.0 | 8.93 | 9.25 | U | AQ |
| *Geochelone gigantea* | 200 | 298 | 15 | 33 | 175.0 | 9.14 | 9.48 | U | AQ |
| *Gopherus agassizii* | 4.500 | 299 | 22 | 47 | 80.0 | 8.97 | 9.30 | Ch,U | AQ |
| *Sphenodon punctatus* | 0.800 | 293 | 18 | 43 | 77.0 | 8.86 | 9.24 | Ch,U | AQ |
| *Crocodylus niloticus* | 400 | 303 | 25 | 49 | 70.0 | 8.96 | 9.26 | Ch,U | AQ |
| *Alligator mississippiensis* | 250 | 302 | 28 | 57 | 50.0 | 8.87 | 9.18 | Ch,U | AQ |
| Raw mean $\bar{\ell}^{\text{raw}}$ | | | | | | | $8.615 \pm 0.290$ | | |
| Corrected mean $\bar{\ell}^{\text{corr}}$ (used in analyses) | | | | | | | $8.929 \pm 0.301$ | | |

**Extended Data Table 8.** Amphibians — Arrhenius-corrected (*n* = 9) Correction method identical to reptiles (Extended Data Table 7). Heart rates from published field recordings at listed *T*_field_; lifespans from AnAge build 15 [24].

| Species | $M$ (kg) | $T_{\text{field}}$ (K) | $f_H^{\text{raw}}$ | $f_H^{\text{corr}}$ | $L$ (yr) | $\ell^{\text{raw}}$ | $\ell^{\text{corr}}$ | Source | Corr. |
| --- | --- | --- | --- | --- | --- | --- | --- | --- | --- |
| *Rana temporaria* | 0.025 | 294 | 25 | 55 | 16.0 | 8.32 | 8.67 | A,Ch | AQ |
| *Rana catesbeiana* | 0.500 | 296 | 20 | 43 | 16.0 | 8.23 | 8.56 | A,Ch | AQ |
| *Bufo bufo* | 0.150 | 293 | 22 | 53 | 36.0 | 8.62 | 9.00 | A,Ch | AQ |
| *Xenopus laevis* | 0.200 | 295 | 20 | 45 | 30.0 | 8.50 | 8.85 | A | AQ |
| *Ambystoma mexicanum* | 0.300 | 294 | 18 | 41 | 25.0 | 8.37 | 8.73 | A | AQ |
| *Salamandra salamandra* | 0.080 | 290 | 20 | 49 | 24.0 | 8.40 | 8.79 | A,Ch | AQ |
| *Plethodon glutinosus* | 0.012 | 291 | 30 | 74 | 20.0 | 8.50 | 8.89 | A | AQ |
| *Necturus maculosus* | 0.130 | 288 | 18 | 46 | 30.0 | 8.45 | 8.86 | A | AQ |
| *Cryptobranchus alleganiensis* | 0.600 | 289 | 15 | 39 | 55.0 | 8.64 | 9.05 | A | AQ |
| Raw mean $\bar{\ell}^{\text{raw}}$ | | | | | | | $8.448 \pm 0.127$ | | |
| Corrected mean $\bar{\ell}^{\text{corr}}$ | | | | | | | $8.822 \pm 0.146$ | | |

#### Dataset summary

Table 14 gives the species counts, body-mass ranges, and ℓ statistics for all eight groups. The complete dataset is provided in Extended Data Tables 1–8 of this paper; no external repository exists. A tab-delimited file is available from the corresponding author on request.

**Table 14:** Summary of the 230-species PBTE dataset. *n*: number of species. *M* : body-mass range (kg). 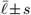: mean s.d. of 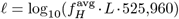 . Δℓ: deviation from the non-primate placental baseline 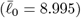.

| Group | $n$ | $M$ range (kg) | $\bar{\ell} \pm s$ | $\Delta\ell$ |
| --- | --- | --- | --- | --- |
| Non-primate placentals | 46 | 0.002–4,000 | $8.998 \pm 0.160$ | 0 (reference) |
| Marsupials / monotremes | 19 | 0.018–30 | $8.933 \pm 0.204$ | −0.062 |
| Primates | 18 | 0.35–160 | $9.376 \pm 0.125$ | +0.381*** |
| Bats | 31 | 0.005–1.1 | $9.540 \pm 0.163$ | +0.545*** |
| Cetaceans (dive-corrected) | 12 | 55–140,000 | $8.801 \pm 0.296$ | −0.194 |
| Birds | 78 | 0.004–115 | $9.528 \pm 0.213$ | +0.533*** |
| Reptiles (Arrhenius-corrected) | 17 | 0.006–400 | $8.929 \pm 0.301$ | −0.065 |
| Amphibians (Arrhenius-corrected) | 9 | 0.012–0.60 | $8.822 \pm 0.146$ | −0.173 |
| **All endotherms** | **194** | | $9.509 \pm 0.397$ | |
| **Full dataset** | **230** | | $9.420 \pm 0.428$ | |

Significance vs non-primate baseline: \* $p < 0.05$ , \*\*\* $p < 0.001$ (Welch $t$ -test).

